# Quickomics: exploring omics data in an intuitive, interactive and informative manner

**DOI:** 10.1101/2021.01.19.427296

**Authors:** Benbo Gao, Jing Zhu, Soumya Negi, Xinmin Zhang, Stefka Gyoneva, Fergal Casey, Ru Wei, Baohong Zhang

## Abstract

**Summary:** We developed Quickomics, a feature-rich R Shiny-powered tool to enable biologists to fully explore complex omics statistical analysis results and perform advanced analysis in an easy-to-use interactive interface. It covers a broad range of secondary and tertiary analytical tasks after primary analysis of omics data is completed. Each functional module is equipped with customizable options and generates both interactive and publication-ready plots to uncover biological insights from data. The modular design makes the tool extensible with ease.

**Availability:** Researchers can experience the functionalities with their own data or demo RNA-Seq and proteomics datasets by using the app hosted at http://quickomics.bxgenomics.com and following the tutorial, https://bit.ly/3rXIyhL. The source code under GPLv3 license is provided at https://github.com/interactivereport/Quickomics for local installation.

**Supplementary information:** Supplementary materials are available at https://bit.ly/37HP17g.

## 1 Introduction

Over the last decade, proteomics and RNA-Seq have become the standard experimental approaches for confidently identifying and accurately quantifying thousands of proteins and genes in complex biological systems. Typically, data generated from those high-throughput experiments are analyzed by data analysts and primary results are provided to end-users as gene-by-sample matrices, tables of summary statistics derived from differential expression analysis, and often a small set of static figures showing high-level results. The large scale of such datasets demands a more effective delivery method beyond the tabulated results, to enable full utilization of a given dataset. Several tools allowing interactive exploration and visualization of complex omics data for end-users without programming skills, such as START (Nelson *et al*., 2017), PIVOT (Zhu *et al*., 2018), PaintOmics 3 (Hernández-De-Diego *et al*., 2018), iSEE (Lun *et al*., 2018), iDEP (Ge *et al*., 2018), WllsON (Schultheis *et al*., 2019), IRIS-EDA (Monier *et al*., 2019), DEBrowser (Kucukural *et al*., 2019), Ideal (Marini *et al*., 2020), and BEAVR (Perampalam and Dick, 2020), have recently been developed. However, these tools come with limitations regarding input formats, the ability to easily adjust plotting parameters, narrow focus on RNA-Seq data analysis, or lack of comprehensive functionalities covering major secondary and tertiary analytical tasks such as gene set enrichment, co-expression network analysis and comparative pathway analysis (detailed feature comparison is outlined in supp. table 1).

To address these gaps, we developed Quickomics, an easy-to-use tool for the visualization of omics (mainly RNA-Seq and proteomics) data and statistical analysis results by leveraging newly developed R packages and modern JavaScript plotting libraries to enhance the usability from data quality control to generation of publication-ready figures.

## 2 Methods

### Architecture

Quickomics is built on R Shiny, a web application framework provided as an R package (http://shiny.rstudio.com/). R Shiny allows applications to be deployed on a personal computer using RStudio or hosted on a local or cloud-based Shiny server. As a Shiny app, Quickomics natively integrates with many R packages, such as ComplexHeatmap (Gu *et al*., 2016) to perform all analytical tasks on the server while presenting results in interactive web pages by utilizing web techniques. In addition, Quickomics adopts a modular design to ensure extensibility.

Highly configurable visualization is one of the major strengths of Quickomics. Often, users have diverse requirements on how to display analyzed data to facilitate interpretation and/or to identify data subsets for further analysis. Here, we take advantage of the dynamic nature of R Shiny where users can easily query data on-the-fly to only display selected groups, samples or proteins/genes of interest.

### Integrated Gene Set Query

We developed xGenesets API (Application Programming Interface) to provide functionality of selecting a pre-defined subset of genes/proteins when it is needed in Quickomics as shown in supp. tutorial section 4.1. Detailed implementation and usage of the API is described in section 1.1 of the tutorial.

### Interactive and High-Resolution Plots

R Shiny controlled input widgets are used to enable users to configure many plotting parameters like font size, color palette, width and height. Moreover, the svglite package was deployed to allow exporting of interactively generated high-resolution plots in PDF or SVG format.

## 3 Results

Quickomics provides a comprehensive visualization workflow for major secondary and tertiary analytical tasks of high dimensional data, which is composed of nine main modules, namely *QC Plots, Volcano Plot, Heatmap, Expression Plot, Gene Set Enrichment, Pattern Clustering, Correlation Network, Venn Diagram* and *Venn Across Projects*, for result visualization and/or analysis, and one *Output* module (details of 37 functions in modules are described in supp. table 2). These modules work essentially the same way for both RNA-seq and proteomics datasets. The detailed guidance about how to prepare, upload and explore a dataset is provided in the supp. tutorial. The main functionalities of Quickomics are exemplified in Fig. 1 by applying the tool to an RNA-seq study (Gyoneva *et al*., 2019), which is available in the demo app along with two proteomics datasets (Connor-Robson *et al*., 2019; Ping *et al*., 2018).

**Fig. 1.**
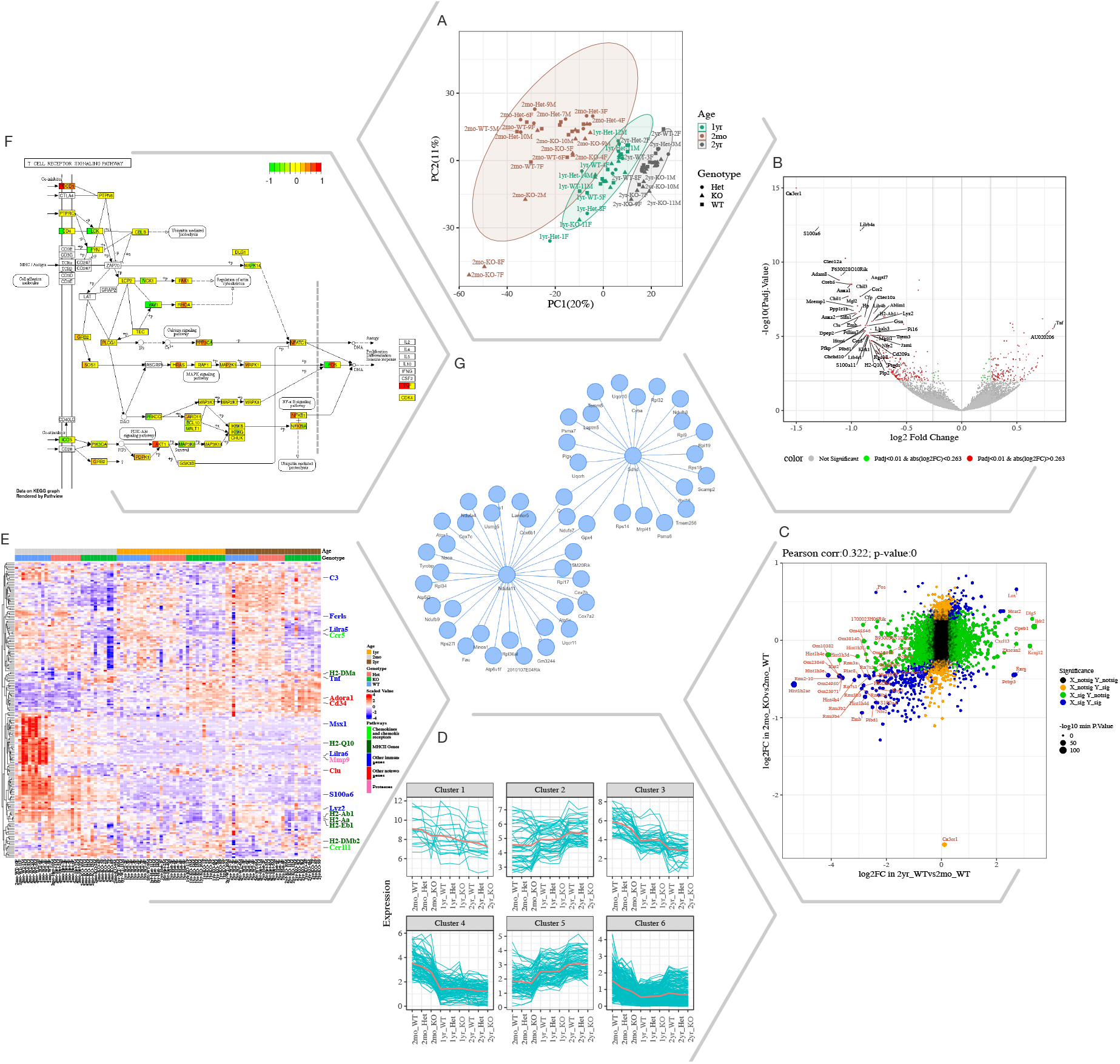
Selected Quickomics functions applied to a dataset of microglial RNA-seq gene expression from three mouse genotypes over time. A) PCA based on full dataset highlights primary sample separation by mouse age at which the cells were isolated. (B) Volcano plot visualizes differentially expressed genes, most of which show reduced expression in 2mo_KO compared to 2mo_WT microglia. For spacing purpose, absolute log2FC (Fold Change) and negative log10 adjusted p-value are capped at 1.5 and 15, respectively. (C) Correlation analysis between two comparisons shows that aging and Cx3cr1-KO have a similar effect on gene expression. (D) Pattern clustering identifies subsets of genes with similar expression over the samples. The clustering is mostly driven by age, with the KO genotype having a similar, but smaller effect. (E) Heatmap of all samples allows the identification of gene clusters with expression regulated by age and/or genotype. Key genes and the pathways they belong to are highlighted on the right. (F) After pathway enrichment analysis, KEGG pathways (Kanehisa and Goto, 2000) of interest can be displayed in a cellular context. The color bars with each stripe representing one comparison show log2 fold changes in various comparisons, allowing project-wide insights for patterns of expression. (G) Correlation network shows potential links between genes of interest. An interactive version of the figure for enlarged view of individual panels is available at https://bit.ly/3psK9tp.

By means of the data uploading tool, Quickomics can ingest datasets consisting of RNA and/or protein expression values (normalized and logarithmic transformed counts or intensity or ratios), statistical measures of comparisons between groups (fold changes, p-values and adjusted p-values), sample tables including sample group and comparison information, and a table containing protein/gene identifiers in text format (supp. tutorial section 2.1). Anticipating differences in naming genes and proteins, a separated section for preparing proteomic dataset is provided in the supp. tutorial section 2.2.2.

Upon loading a dataset, users can perform a comprehensive Quality Control (QC) analysis by checking the expression similarity of samples within and across experimental groups using principal component analysis (PCA), a box-and-whisker plot, a coefficient of variation (CV) distribution plot and a sample-sample distance matrix. PCA allows users to detect outliers or batch effects across groups and/or samples, which can be viewed in 2D, 3D, and 3D Interactive plots, and to compare the similarity of experimental groups to one another. The heatmap and dendrograms visualize the similarity of all the samples. The box plot and CV distribution facilitate assessment of quantitative precision across samples quantification and mean variation distribution, respectively. These plots can be dynamically re-plotted upon user selection of a subset of groups and/or samples.

Then, users can visualize genes/proteins showing different expression patterns across groups via *Volcano Plot* and *Expression Plot* and identify correlated genes/proteins from *Pattern Clustering* and *Correlation Network* modules.

The *Volcano plot* is commonly used to visualize immediate outliers, i.e. significantly differentiated proteins/genes with large fold change (FC), in two-group comparisons by both static and interactive volcano plots (log2 FC versus negative log10 P value). It could also be used for a quick assessment of the fraction of the altered proteome or genome. This module also provides an innovative function for users to visualize the expression changes for differentially expressed genes between two comparisons as shown in Fig.1C.

The *Heatmap* provides both static and interactive gene/protein expression heatmap visualization options to facilitate discovery of similarity patterns across samples based on user-defined parameters. Users can define sample group(s) and gene sets to be visualized and also customize the clustering options.

The *Expression Plot* allows users to visualize the expression of proteins/genes that passed the fold change and P value cutoffs in group comparisons with multiple plot options. The users can also search for genes and visualize expression data in a graphical or tabular format. The *Pattern Clustering* provides clustering analysis based on user defined interesting gene/protein subsets to visualize the gene/protein expression patterns across different groups (e.g., time points/different conditions). Several clustering methods including soft clustering, k-means and partitioning around medoids are available. The *Correlation Network* generates a co-expression network constructed based on protein-protein/gene-gene correlation matrix. In order to speed up the response time, only correlation pairs with r2 > 0.7 were used.

After expression analysis, users can perform a functional enrichment analysis on differentially expressed proteins/genes from a selected comparison against pathways and gene sets to get biological insights using the *Gene Set Enrichment* module. After enrichment analysis, the expression value of query proteins/genes is listed in a table and can be visualized in heatmap from all the studied groups in the whole dataset. If the functional pathways used in the gene set enrichment analysis are from KEGG, a KEGG pathway graph will be available with query genes overlaid by log2 fold change values from one or more comparisons.

Furthermore, *Venn Diagram* modules were incorporated to enable data comparison within and across projects. The *Venn Diagram* allows users to generate Venn diagram among different comparisons in the same experiment. The *Venn Diagram Across* Projects allows users to generate Venn diagram among different experiments uploaded by users. Overlapping genes/proteins are available in Intersection Output section.

Quickomics generates highly customized plots and further provides interactive forms of PCA 3D plot, volcano plot, heatmap, and correlation network, which allows users to hover mouse over individual data points to view annotation information. In the end, Quickomics contains an *Output* module to export selected figures in high resolution suitable for publication as well as download data tables in excel format.

## 4 Conclusions

In summary, Quickomics is a powerful tool to help biologists explore complex omics data and interpret results through a user-friendly interface. Firstly, it provides comprehensive quality control analysis with adjustable options for visualization. Secondly, it supports most of the major secondary and tertiary analytical tasks including volcano plot, heatmap, expression plot, gene set enrichment, pattern clustering, correlation network and Venn diagram. Finally, it is released as open source to promote suggestions for new features and contributions from the bioinformatics community to further enhance this versatile tool.

## Supporting information

Supplemental Table 1, 2 and tutorial

