## Supplemental Table 1, 2 and tutorial for "Quickomics: exploring omics data in an intuitive, interactive and informative manner"

Table S1. Comparison of features of omics data visualization tools. The table in excel format is available at https://bit.ly/3qax8VZ for better view.

| Tools | Quickomics | Ideal | BEAVR | WIlsON | IRIS-EDA | DEBrowser | PIVOT | PaintOmics 3 | iSEE | IDEP | START |
|---|---|---|---|---|---|---|---|---|---|---|---|
| **Year** | 2020 | 2020 | 2020 | 2019 | 2019 | 2019 | 2018 | 2018 | 2018 | 2018 | 2017 |
| **Main function** | | | | | | | | | | | |
| Data analysis | Y | Y | Y | | Y | Y | Y | | | Y | Y |
| Data visualization | Y | Y | Y | Y | Y | Y | Y | Y | Y | Y | Y |
| **Data type supported** | | | | | | | | | | | |
| RNAseq | Y | Y | Y | Y | Y | Y | Y | Y | Y | Y | Y |
| Other datatypes | Proteomics | | | Multi-omics | | Other sequencing | | Multi-omics | Multi-omics | | |
| **Key features** | Comprehensive analytical tasks, generates both interactive and publication-ready figures. | Interactive differential expression analysis for RNAseq. Generate publication-ready outputs. | Simplifies RNAseq analysis for novice users/experts | Provides customized and dynamic visualization for multi-omics data | Performs RNAseq data analysis and provides a framework to expedite data submission to NCBI's Gene Expression Omnibus | Interactive differential expression analysis and visualization for count data | Interactive analysis and visualization of transcriptomics data, automatic report generation, publication-quality plots. | Integrated visualization of multiple omic data types onto KEGG pathway diagrams. | Provides a general visual interface for exploring data in a SummarizedExperiment object. | Integrated web application for differential expression and pathway analysis of RNA-Seq data | Web-based RNAseq analysis and visualization |
| **Online/Standalone** | Both | Both | Standalone | Standalone | Online | Standalone | Standalone | Online | Standalone | Both | Both |
| **Source Code Link** | https://github.com/interactivereport/Quickomics | https://github.com/federicomarini/ideal | https://github.com/developerpiru/BEAVR | https://github.molgen.mpg.de/loosolab/wilson-apps | https://github.com/OSU-BMBL/IRIS | https://github.com/UMMS-Biocore/debrowser | https://github.com/qinzhu/PIVOT | https://github.com/fikipollo/paintomics3 | https://github.com/iSEE/iSEE | https://github.com/iDEP-SDSU/idep | https://github.com/jminnier/STARTapp |
| **Demo Link** | http://quickomics.bxgenomics.com/ | http://shiny.imbei.uni-mainz.de:3838/ideal/ | | http://loosolab.mpi-bn.mpg.de/wilson/ | http://bmbl.sdstate.edu/IRIS/ | | https://kim.bio.upenn.edu/software/pivot.shtml | http://www.paintomics.org/ | | http://bioinformatics.sdstate.edu/idep/ | https://kcvi.shinyapps.io/START |
| **Demo using example data** | Y | Y | | Y | Y | | | | | Y | Y |
| **Analysis tool** | | | | | | | | | | | |
| PCA-2D | Y | | Y | Y | Y | Y | Y | | Y | Y | Y |
| PCA-3D | Y | | | | | | Y | | | | |
| PCA 3D interactive plot | Y | | | | | | Y | | | | |
| Sample-sample distance heatmap | Y | Y | Y | Y | Y | | Y | | | | Y |
| Sample to sample scatter plots | | Y | | | Y | Y | Y | | | Y | |
| Dendrograms | Y | | Y | Y | Y | | Y | | | Y | |
| Rank-frequency plot/Mean-Variability Plot/Rank-Sd plot | | | | | | | Y | | | | |
| Gene expression per sample in box plot | Y | | | | Y | Y | | | | Y | |
| CV Distribution plot | Y | | | | | Y | Y | | | | |
| Multidimensional scaling plot | | | | | Y | | Y | | | Y | |
| t-distributed Stochastic Neighbor Embedding plot | | | | | Y | | Y | | Y | | |
| Histogram for read counts by sample/Bar plot of total reads by sample | | | | | Y | Y | Y | | | Y | |
| Differential expression analysis | | Y | Y | | Y | Y | Y | | | Y | |
| Bar plot for DEG number overview | | | | | Y | | | | | | |
| Static volcano plot | Y | Y | Y | Y | Y | Y | | | | Y | |
| Interactive volcano plot | Y | | | | Y | | | | | | Y |
| MA plots | | Y | | Y | Y | Y | | | | Y | |
| Scatter plot | | | | | | Y | | | Y | | Y |
| Rectangle plot | | | | | | | | | Y | | |
| Histograms for unadjusted p-values/histogram for estimated log fold changes/Schweder-Spjøtvoll plot | | Y | | | | | | | | | |
| Fold change vs fold change from two comparisions plot | Y | | | Y | | | | | | | |
| Static gene expression heatmap | Y | Y | Y | Y | Y | Y | Y | Y | Y | Y | Y |
| Interactive gene expression heatmap | Y | Y | | Y | Y | | | | | | Y |
| Gene expression box plot | Y | Y | Y | Y | | | | | | | Y |
| Gene expression jitter plot | | | Y | | | | | | | | |
| Gene expression bar plot | Y | | | Y | | Y | | | | Y | |
| Gene expression violin plot | Y | | | Y | | | Y | | Y | | |
| Gene expression line plot | Y | | | Y | | | | | | | |
| Rank Abundance Curve | Y | | | | | | | | | | |
| Minimum Spanning Tree and Community Detection | | | | | | | Y | | | | |
| Funtional enrichment analysis | Y | Y | Y | | Y | Y | Y | Y | | Y | |
| Interactive pathway network | | | | | | | | Y | | | |
| Multi-omic visualization of single pathways | | | | | | | | Y | | | |
| Gene expression pattern clustering | Y | | | | Y | | | | | Y | |
| Co-expression network | Y | | | | | | Y | | | Y | |
| Visualizing expression profiles on chromosomes | | | | | | | | | | Y | |
| Venn Diagram for single dataset | Y | | | | | | | | | Y | |
| Venn Diagram across datasets | Y | Y | | | | | | | | | |
| UpSet plots | | Y | | | | | | | | | |
| GEO Submission | | | | | Y | | | | | | |

Table S2. Available modules and analysis in Quickomics.

| Modules/Analysis | Description |
|---|---|
| **QC Plots** | |
| PCA Plot | To perform principal component analysis (PCA) and generate a plot that can be colored, shaped and sized by attributes of samples. It has options to emphasize mean points and/or add marginal rugs to the plot. |
| Eigenvalues | To generate a plot of variances explained by the first 10 PCs in the dataset to allow users to make educated decisions of which PCs to plot in 2D or 3D scatter plot. |
| PCA 3D Plot | To display 3D PCA representation on PC1, PC2 and PC3. Users can zoom in and out to focus on different areas of the plot or rotate the plot in different axes to find the best separation. Users can change color, include an ellipsoid and add labels to the samples. |
| PCA 3D Interactive | To create interactive 3D PCA plot to enable users to see detailed sample annotation information by hovering over samples on the plot. It allows users to change color and shape of the dots representing samples in the plot. |
| Sample-sample Distance | To generate a distance matrix for every sample pair and plot it as a heatmap to show pair-wise similarity between all samples. The rows and columns are clustered based on similarity scores. |
| Dendrogram | To generate a Dendrogram plot to help visualize hierarchical clustering relationships between samples. The default plot is circular and divided into four parts. Users have the option to visualize the dendrogram plot in tree horizontally or vertically and divide the plot in multiple regions. |
| Box Plot | To generate a plot to visualize the distribution of the normalized expression values in all samples. This identifies the minimum, first quartile, median, third quartile, and maximum values in the dataset. |
| CV Distribution | To generate a plot to show the histogram of coefficient of variation (CV) and a dotted line for each group indicating the median CV. |
| Order Groups | To allow users to select and/or re-order the groups. The order will be stored in current session and will be displayed in a preferred order in all plots. Users can either drag-n-drop group names to reorder or use the select panel to delete and add. |
| **Volcano Plot** | |
| Volcano Plot (Statics) | To generate a volcano plot to visualize the differentially expressed genes/proteins provided in the input data file. Users have the option to change cutoffs of fold change and P value and more advanced display options. |
| Volcano Plot (Interactive) | To create an interactive volcano plot allowing users to see detailed annotation information of genes/proteins (dots in plot) by hovering over the dots on the plot. |

| | |
|---|---|
| DEGs in Two Comparisons | To generate a scatter plot to visualize the relationship between differentially expressed genes/proteins in two comparisons. |
| Data Table | To view the differentially expressed genes/proteins in a tabular format with a searchable feature. Users could download the table as a CSV file. |
| **Heatmap** | |
| Static Heatmap Layout 1 | To generate a heatmap of gene/protein-by-sample gene expression matrix by using ComplexHeatmap package. Users can customize the gene list and pick annotation categories to be shown on the heatmap. |
| Static Heatmap Layout 2 | To generate a heatmap using the heatmap.2 function from gplots package. The same customization of the above function is applied here. |
| Interactive Heatmap | To generate an interactive heatmap of gene/protein-by-sample gene expression matrix by using package 'heatmaply'. |
| **Expression Plot** | |
| Browsing | To generate Violin Plots to visualize the expression of the differentially expressed genes/proteins identified in various comparisons. |
| Searched Expression Data | To generate Box plots to visualize the expression of user entered genes/proteins list or user selected genes/proteins list from databases like KEGG, MSigDB etc. provided in Quickomics. |
| Data Table | A searchable table for the genes/proteins selected in "Searched Expression Data" tab with their normalized expression values. |
| Result Table | A searchable table for the genes/proteins selected in "Searched Expression Data" tab with their statistics results including P value, adjusted P value and fold change. |
| Rank Abundance Curve | To generate a plot to visualize the relative abundance of a list of genes within a dataset. This plot helps interpret the distribution of abundance and expression levels of a set of genes. |
| **Gene Set Enrichment** | |
| Gene Set Enrichment | To perform gene set enrichment analysis based on user selected gene set and functional databases. |
| Gene Expression | To provide logarithm transformed fold change values for genes in the enriched functional pathways identified from the above analysis. |
| Gene Set Heatmap | To generate expression heatmap for genes in the enriched functional pathways identified in the "Gene Set Enrichment" analysis. |
| KEGG Pathway View | To allow users to view the fold change levels of genes in the KEGG pathways for up to 5 comparisons together. |
| **Pattern Clustering** | |

| | |
|---|---|
| Clustering of Centroid Profiles | To perform genes/proteins clustering based on their expression values across different groups and generate plots to visualize the identified co-expression clusters. Three clustering algorithms including soft (fuzzy) clustering, k-means and partitioning around medoids are available. |
| Data Table | A table to show the expression values for genes/proteins with their co-expression cluster IDs assigned in the "Clustering of Centroid Profiles" section. |
| **Correlation Network** | |
| visNetwork | To build co-expression networks based on gene-gene or protein-protein correlation using R package visNetwork. Users can choose the correlation coefficient and P value cutoffs to select expression correlated genes/proteins. |
| networkD3 | To build co-expression networks based on gene-gene or protein-protein correlation using R package networkD3. Users could drag, zoom, and highlight gene/protein nodes in the network. |
| Data Table | A table to show genes/proteins in the co-expression networks identified in the "visNetwork" section. Users can perform searches and sort the table based on correlation statistics. |
| **Venn Diagram** | |
| Venn Diagram | To generate a Venn Diagram to show the intersections of differentially expressed genes/proteins from up to 5 different comparisons. Users can use all differentially expressed genes/proteins or only up- or down-regulated ones. |
| Venn Diagram (black & white) | To generate a black and white Venn Diagram to show the intersections of differentially expressed genes/proteins from up to 5 different comparisons. |
| Intersection Output | Gene lists from different intersecting regions of the Venn Diagram generated in the "Venn Diagram" section. |
| DEG Table | A table to show fold changes and P values of genes from the Venn Diagram in the comparisons being queried. |
| **Venn Across Projects** | |
| Venn Diagram | To generate a Venn Diagram to show the intersections of differentially expressed genes/proteins across projects. This function could be used to identify common differentially expressed genes/proteins present in RNAseq and Proteomics datasets generated from the same samples. |
| Venn Diagram (black & white) | To generate a black and white Venn Diagram to show the intersections of differentially expressed genes/proteins across projects. |
| Intersection Output | Gene lists from different intersecting regions of the Venn Diagram generated in the "Venn Diagram" section. |

### Quickomics Supplementary Tutorial

### 1 Introduction

The Quickomics tool can be accessed via the link http://quickomics.bxgenomics.com. Implemented through R Shiny, it helps with visualizing statistical analysis results for RNAseq and Proteomics datasets. This supplemental tutorial provides a detailed guide on using the different functionalities and customizing the tools to best fit individual analysis and plotting needs, using published RNAseq dataset and proteomics dataset as examples (Gyoneva *et al.*, 2019; Connor-Robson *et al.*, 2019).

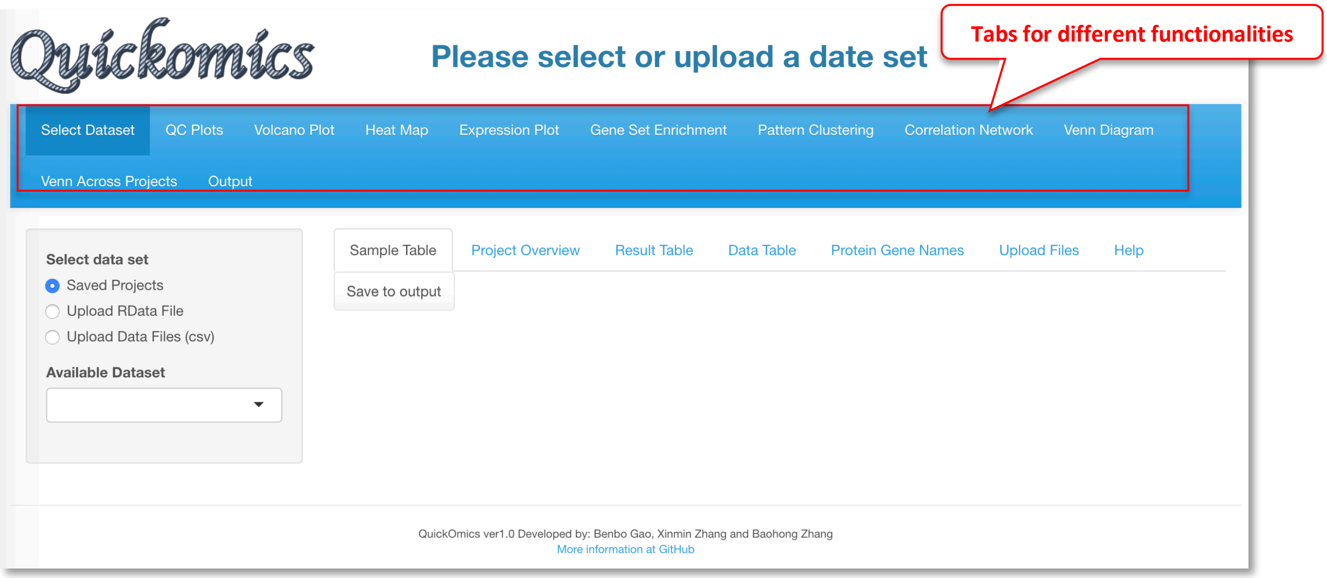

The interface contains multiple tabs, corresponding to different functional modules, that can be accessed on top panel of the webpage. Users have the option to upload their own dataset or choose from existing demo datasets for visualization.

#### 1.1 Integrated Gene Set Query

To supply genes in pre-defined gene sets to plotting functions in Quickomics, we developed xGenesets API for Quickomics to query and retrieve a list of genes/proteins when it is needed. The lists can be defined based on pathways, and gene sets from various sources, including KEGG Pathways (Kanehisa and Goto, 2000), WikiPathways (Martens *et al.*, 2020), Small Molecule Pathway Database (Frolkis *et al.*, 2010), Reactome (Jassal *et al.*, 2020), Gene Ontology (The Gene Ontology Consortium, 2019), Molecular Signatures (Subramanian *et al.*, 2005; Liberzon *et al.*, 2015), and LIPID MAPS Proteome Database (Cotter *et al.*, 2006). xGenesets database currently has 175,537 gene sets stored in MySQL tables, covering human (115,497 records), mouse (30,554 records) and rat (29,486 records) gene sets. xGenesets features personalized settings, dynamic drop-down list upon typing for quick gene set selection, full gene set database browsing, and a list of genes from any gene sets. Throughout Quickomics, users interact with this convenient tool in appropriate interface when a gene set is needed by checking "Geneset" option and then clicking on "Select Geneset" to pick a set from a table of available gene sets in a popup window as shown in the following screenshot.

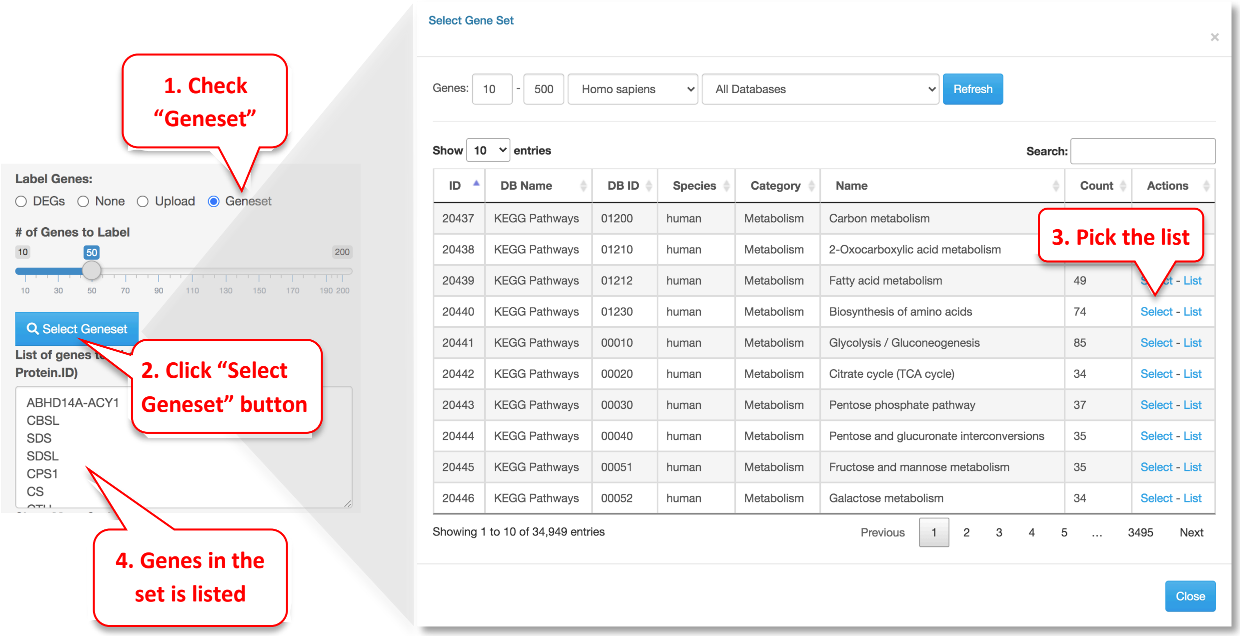

For developers of web-based applications (general user can skip this paragraph), xGenesets can be easily embedded in any modern websites and online applications, e.g., R Shiny web apps, by leveraging jQuery, DataTables, and bootstrap (either v3 or v4) JavaScript libraries. xGenesets API is freely available for all usages. Examples of using xGenesets API and guidance on setting up the API in your own applications can be found at [https://bxaf.net/genesets](https://bxaf.net/genesets).

### 2 Select Dataset Module

This first module is for selecting dataset. Users can either select from pre-loaded example datasets or upload their own dataset in a pre-defined format as detailed in section 2.1 and in our GitHub page (https://github.com/interactivereport/Quickomics). We have pre-loaded four datasets from published studies that users can use for demonstrative purposes. These includes two RNAseq datasets from Gyoneva *et al.*, 2019 and Connor-Robson *et al.*, 2019; and two proteomics datasets from Connor-Robson *et al.*, 2019. Users can walk through multiple sub-tabs to visualize, select and download figures or data.

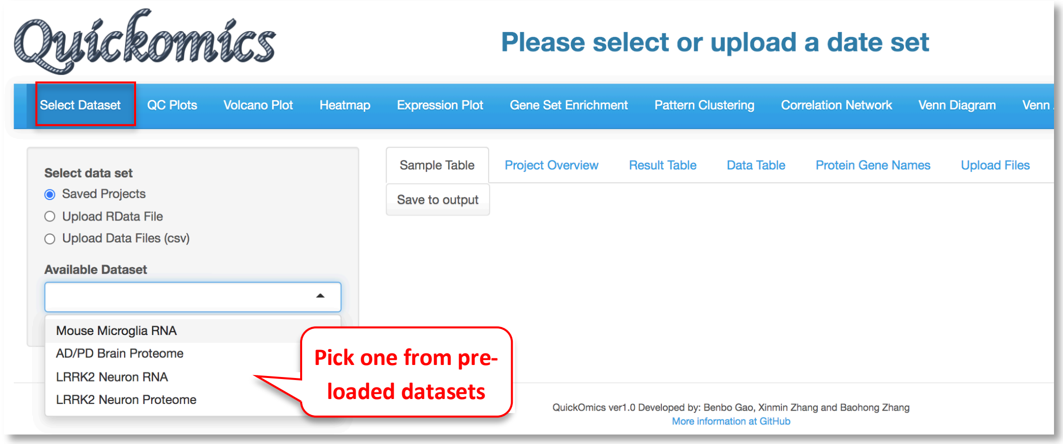

#### 2.1 Upload Files

For a data set, the "Upload Files" tool allows users to upload three required files, namely sample metadata, normalized expression data and statistical comparison results in csv format (Comma Separated Values) to Quickomics directly. Example data sets are provided in GitHub for both RNAseq (https://bit.ly/2MRkFcb) and proteomics (https://bit.ly/3rn4i6a). Detailed formatting guidance is outlined below,

1. **Sample Metadata File:** It should have “sampleid” and “group” columns, with additional columns optional. Sample identifiers must match those used in the expression data file.
2. **Expression Data File:** It should be a matrix of expression values with genes/proteins as rows, and samples as columns. The unique IDs for genes/proteins are in the first column. We recommend using log of normalized expression values, e.g. log2(TPM+1) for RNAseq data or normalized intensity or ratio for proteomics data.
3. **Comparison Data File:** It should have five columns, “UniqueID”, “test”, “Adj.P.Value”, “P.Value” and “logFC”. The comparison names are listed in “test” column. Please note that wrongly named column headers will cause issues.
4. **Optional Gene/Protein Name File:** The system has built-in function to convert unique IDs in the data files to gene symbols and create the Gene/Protein Name file, so most users don't need to prepare the file. Nevertheless, if provided by users, it must have four columns: “id” (sequential numbers like 1,2,3 … …), “UniqueID” (matching IDs used in the expression and comparison data file), “Gene.Name” (official gene symbols), “Protein.ID” (UniProt protein IDs, or keep it empty for RNA-Seq data). Additional columns (e.g. gene biotype) are optional.

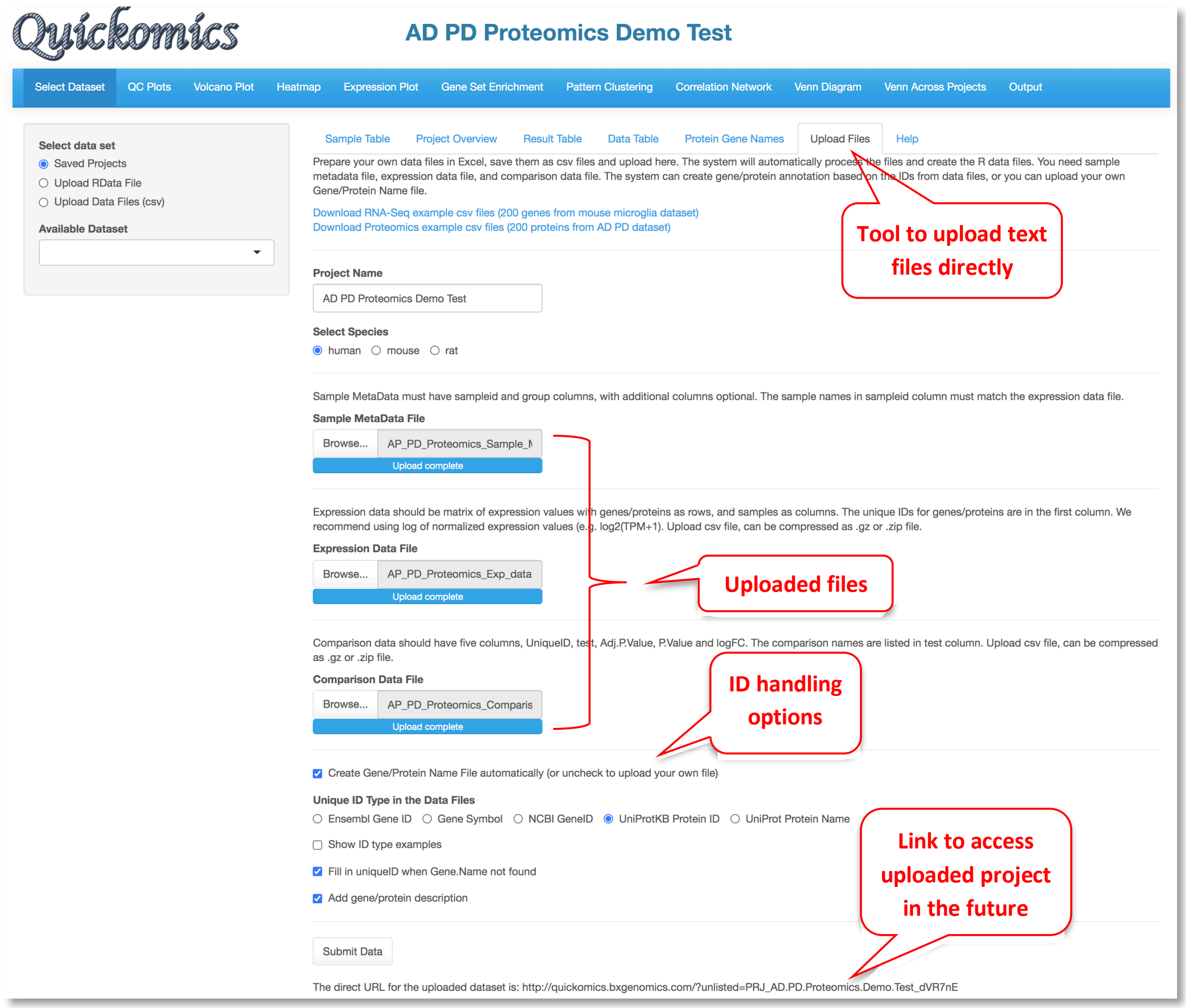

After the data files are processed, Quickomics will automatically load all required data for exploration immediately and provide a link for the user to come back in the future.

Behind the scene, Bioconductor biomaRt package (https://bioconductor.org/packages/release/bioc/html/biomaRt.html) has been used to convert gene IDs (Ensembl gene, NCBI gene ID, etc.) into gene symbols by querying Ensembl databases. For protein IDs, we generated a custom lookup table using information downloaded from UniProt Knowledgebase to convert UniProt IDs to gene symbols and protein names. We didn't use biomaRt for proteins as Ensembl databases only cover about 60-80% protein IDs in a typical proteomics data set.

#### 2.2 Prepare R Data Files by Computational Biologists

We recommend uploading csv files, which is convenient for general users who can skip section 2.2 entirely. Nevertheless, experienced R programmers can create R data files to be uploaded through "Upload RData File" option. Section 2.2.1 and 2.2.2 provide example R scripts to prepare such R data files for RNA-Seq and proteomics data, respectively.

Two R data files are required for each data set, one contains the main data and the other contains gene co-expression network information. For the pre-loaded datasets, main data files are located in the "data" folder, https://github.com/interactivereport/Quickomics/tree/master/data, and gene co-expression network files are located in the "networkdata" folder, https://github.com/interactivereport/Quickomics/tree/master/networkdata. One can review the content of a R data file (e.g. Mouse_microglia_RNA-Seq.RData) in the "data" folder by loading it into R. The main R data file contains the following R data frame objects.

1. **MetaData**: It must have "sampleid", "group", "Order" and "ComparePairs" columns. Additional metadata columns about samples are optional. "sampleid" should match those used in expression data. "group" holds group names of samples. "Order" is ordered group names used on plotting. "ComparePairs" are names of comparisons performed.
2. **ProteinGeneName**: It must have "UniqueID", "Gene.Name" and "Protein.ID" columns. "UniqueID" matches gene ID in below data_wide and data_long objects. "Gene.Name" should be official gene symbols. "Protein.ID" is UniProt protein IDs, or empty for RNA-Seq data. Additional columns about proteins or genes are optional.
3. **data_wide**: This is the expression matrix in which rows are genes and columns are samples. Samples must match "sampleid" values in MetaData and gene IDs must match "UniqueID" values in ProteinGeneName.
4. **data_long**: Gene expression matrix in long format with four columns, "UniqueID", "sampleid", "expr" and "group". "group" values must match those listed in MetaData.
5. **results_long**: The comparison results in long format with five columns, "UniqueID", "test", "Adj.P.Value", "P.Value" and "logFC". "UniqueID" matches "UniqueID" in ProteinGeneName. "test" column has the comparison names that must match "ComparePairs" values in MetaData. The other values are typically computed from statistical analysis, but the data headers must be changed to "Adj.P.Value", "P.Value" and "logFC".
6. **data_results**: This is a summary table starting with "UniqueID" and "Gene.Name" columns, then the intensity (max or mean expression value from data_wide for each gene), mean and SD expression values for each group, and finally comparison data (comparison name added as prefix of columns).

The network data object is computed from "data_wide" expression matrix by using Hmisc R package exemplified by the code snippet below.

```
cor_res <- Hmisc::rcorr(as.matrix(t(data_wide)))
cormat <- cor_res$r
pmat <- cor_res$P
ut <- upper.tri(cormat)
network <- tibble::tibble (
  from = rownames(cormat)[row(cormat)[ut]],
  to = rownames(cormat)[col(cormat)[ut]],
  cor  = signif(cormat[ut], 2),
  p = signif(pmat[ut], 2),
  direction = as.integer(sign(cormat[ut]))
)
```

##### 2.2.1 Example R script to prepare R data files from RNA-Seq results

We have provided example input files (TPM and count matrix files, sample grouping file, comparison list file) and the R scripts to generate the main data and network R data files at https://github.com/interactivereport/Quickomics/tree/master/demo_files/Example_RNA_Seq_data. Please note that you may need to modify RNA_Seq_raw2quickomics.R to fit your input files.

- rsem_TPM.txt: The TPM matrix. One can also use RPKM matrix if needed.
- rsem_expected_count.txt: The gene count matrix. We used RSEM counts in this case, but gene count results from other methods can be used as well.
- grpID.txt: This file lists the group information for each sample.
- comparison.txt: This list lists the comparisons to perform (group 1 vs group 2 in each row).

The following command will read the above data files, run differential gene expression analysis using DESeq2, and create main and network R data files.

```
$ Rscript RNA_Seq_raw2quickomics.R
```

##### 2.2.2 Example R script to prepare R data files from proteomics results

We have provided the example input files (normalized protein expression, comparison data, sample information, protein and gene names) and the R script to generate the main data and network R data files at https://github.com/interactivereport/Quickomics/tree/master/demo_files/Example_Proteomics_data. Please note that you may need to modify Proteomics2Quickomics.R to fit your input files.

- NormalizedExpression.csv: Normalized protein expression (log2 transformed).
- ComparisonData.csv: Comparison results. The statistic values are: logFC, P.Value and Adj.P.Value. This can be created using R packages like limma.
- Sample.csv: Sample information file.
- ProteinID_Symbol.csv: This file lists the proteinIDs and associate gene symbols.

The following command will read the above data files and create main and network R data files.

```
$ Rscript Proteomics2Quickomics.R
```

#### 2.3 Sample Table

A sample table with metadata/details appears upon completion of dataset loading. Here we have selected the "Mouse Microglia RNA" dataset from the previously published paper (Gyoneva *et al.*, 2019) as an example to illustrate all the functionalities. This publication used RNAseq to conclude that the knockout of the Cx3cr1 gene altered the microglial transcriptome in a manner similar to what ageing does, albeit in a milder extent. Using Quickomics, we can visualize the original results from the paper, as well as provide novel visualizations to support new results.

The Sample Table contains detailed info of all study samples. A total of 93 samples that were sequenced in this project are listed in the table with attributes like sampleid, group, Age, Genotype and Gender, shown in different columns.

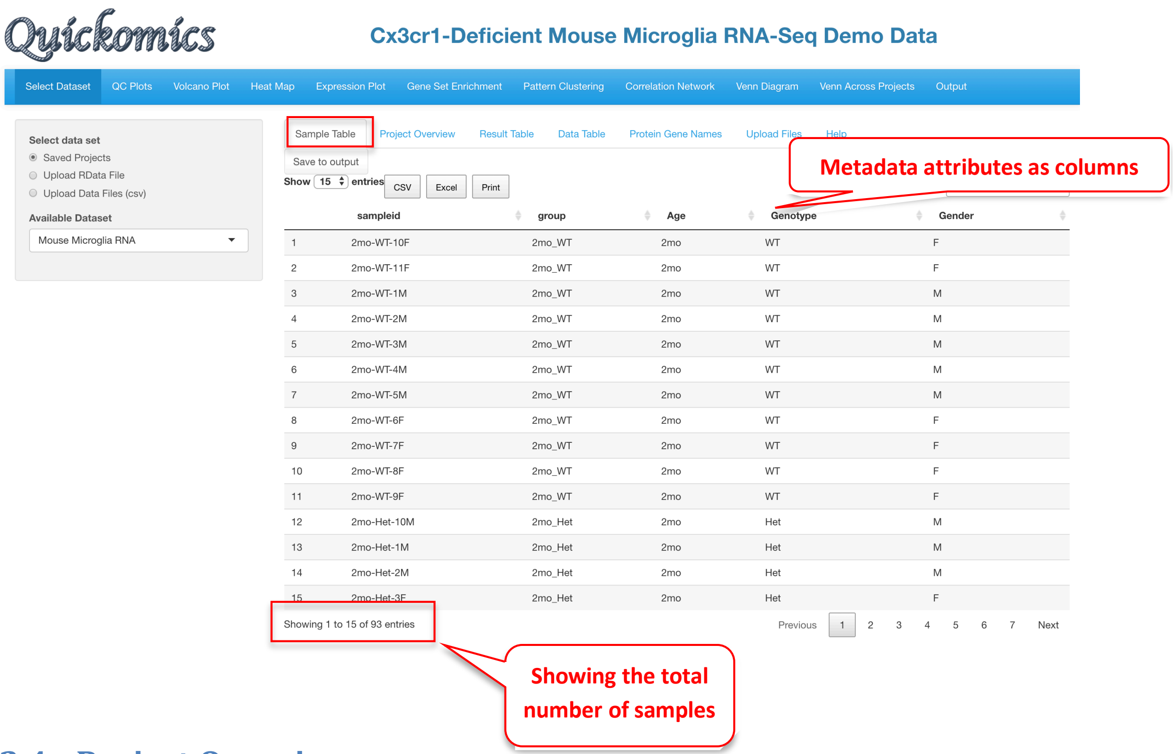

#### 2.4 Project Overview

This tab gives an overview of all samples in the project by providing a summary from the metadata. This includes information like which species was used, the different comparisons run and total number of samples in each group. This tab also helps identify how many groups were present in the dataset, like in this project there are 9 different groups that correspond to three genotypes in each of three time points (WT, Het, KO; 2mo, 1yr, 2yr).

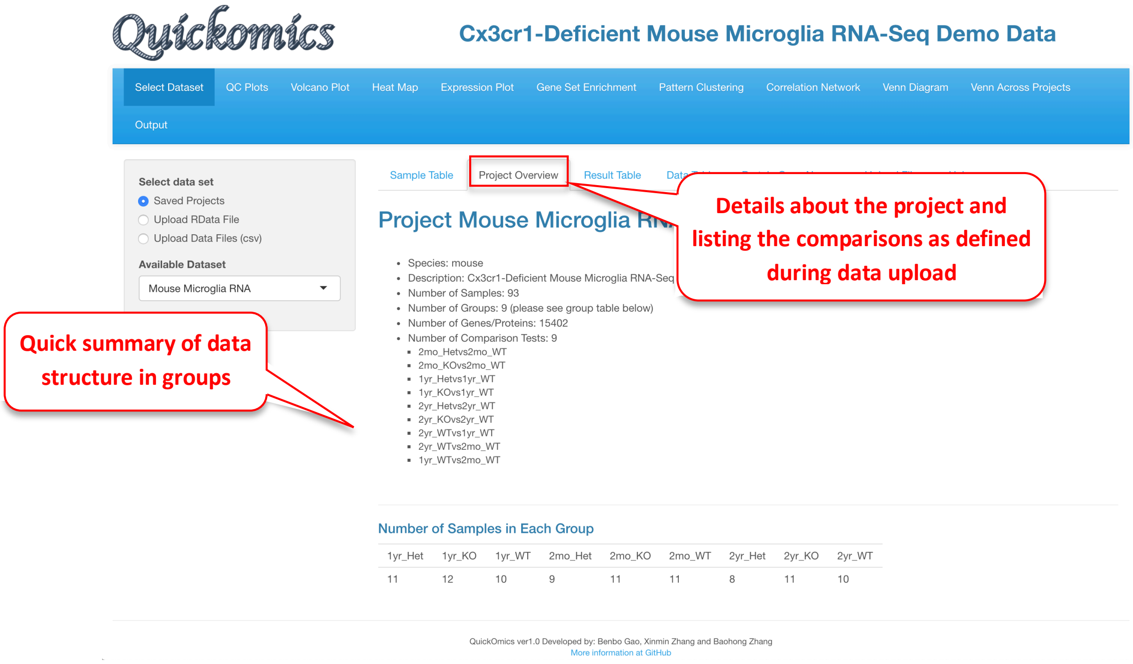

#### 2.5 Result Table

This tab contains statistical analysis results like Ensemble ID (for RNAseq data) and Accession ID (for proteomics data), gene name, mRNA or protein abundance (count or intensity), mean and SD values for each group, log2 fold change, P value and adjusted P value.

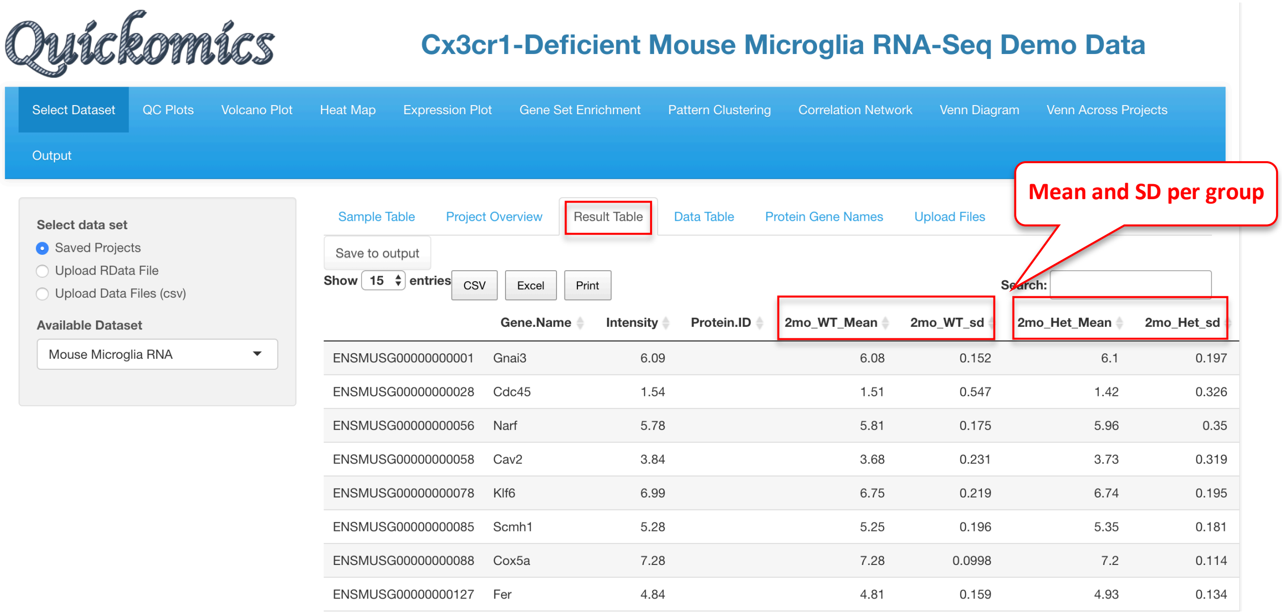

| | Gene.Name | Intensity | Protein.ID | 2mo_WT_Mean | 2mo_WT_sd | 2mo_Het_Mean | 2mo_Het_sd |
|---|---|---|---|---|---|---|---|
| ENSMUSG00000000001 | Gnai3 | 6.09 | | 6.08 | 0.152 | 6.1 | 0.197 |
| ENSMUSG00000000028 | Cdc45 | 1.54 | | 1.51 | 0.547 | 1.42 | 0.326 |
| ENSMUSG00000000056 | Narf | 5.78 | | 5.81 | 0.175 | 5.96 | 0.35 |
| ENSMUSG00000000058 | Cav2 | 3.84 | | 3.68 | 0.231 | 3.73 | 0.319 |
| ENSMUSG00000000078 | Klf6 | 6.99 | | 6.75 | 0.219 | 6.74 | 0.195 |
| ENSMUSG00000000085 | Scmh1 | 5.28 | | 5.25 | 0.196 | 5.35 | 0.181 |
| ENSMUSG00000000088 | Cox5a | 7.28 | | 7.28 | 0.0998 | 7.2 | 0.114 |
| ENSMUSG00000000127 | Fer | 4.84 | | 4.81 | 0.159 | 4.93 | 0.134 |

#### 2.6 Data Table

The Data table contains normalized expression values (TPM for RNAseq, intensity for label free proteomic quantification, or ratio for isobaric label proteomics quantification) per sample for each gene.

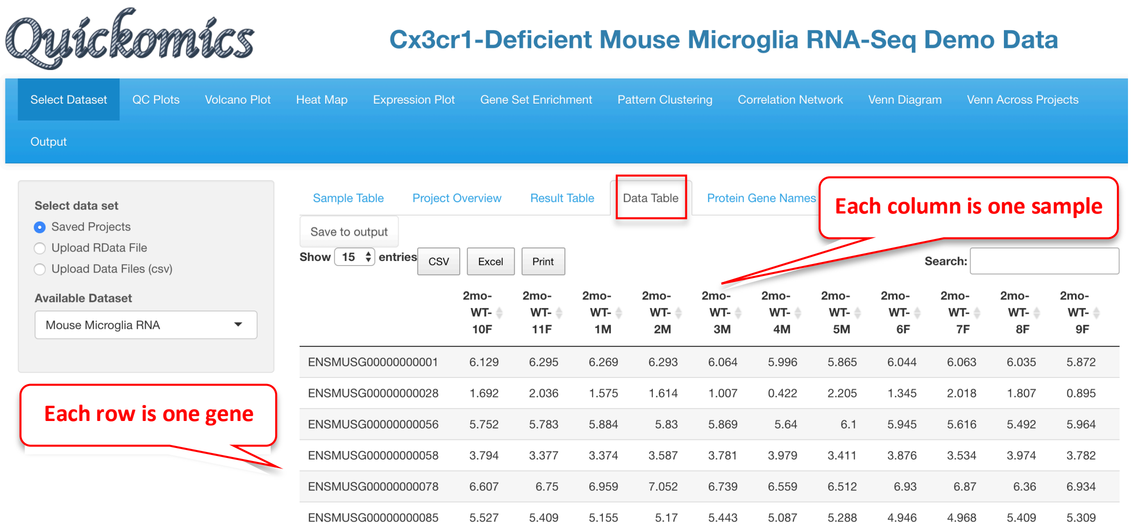

| | 2mo-WT-10F | 2mo-WT-11F | 2mo-WT-1M | 2mo-WT-2M | 2mo-WT-3M | 2mo-WT-4M | 2mo-WT-5M | 2mo-WT-6F | 2mo-WT-7F | 2mo-WT-8F | 2mo-WT-9F |
|---|---|---|---|---|---|---|---|---|---|---|---|
| ENSMUSG00000000001 | 6.129 | 6.295 | 6.269 | 6.293 | 6.064 | 5.996 | 5.865 | 6.044 | 6.063 | 6.035 | 5.872 |
| ENSMUSG00000000028 | 1.692 | 2.036 | 1.575 | 1.614 | 1.007 | 0.422 | 2.205 | 1.345 | 2.018 | 1.807 | 0.895 |
| ENSMUSG00000000056 | 5.752 | 5.783 | 5.884 | 5.83 | 5.869 | 5.64 | 6.1 | 5.945 | 5.616 | 5.492 | 5.964 |
| ENSMUSG00000000058 | 3.794 | 3.377 | 3.374 | 3.587 | 3.781 | 3.979 | 3.411 | 3.876 | 3.534 | 3.974 | 3.782 |
| ENSMUSG00000000078 | 6.607 | 6.75 | 6.959 | 7.052 | 6.739 | 6.559 | 6.512 | 6.93 | 6.87 | 6.36 | 6.934 |
| ENSMUSG00000000085 | 5.527 | 5.409 | 5.155 | 5.17 | 5.443 | 5.087 | 5.288 | 4.946 | 4.968 | 5.409 | 5.309 |

#### 2.7 Protein Gene Names

This tab contains all quantified gene or protein IDs and gene names.

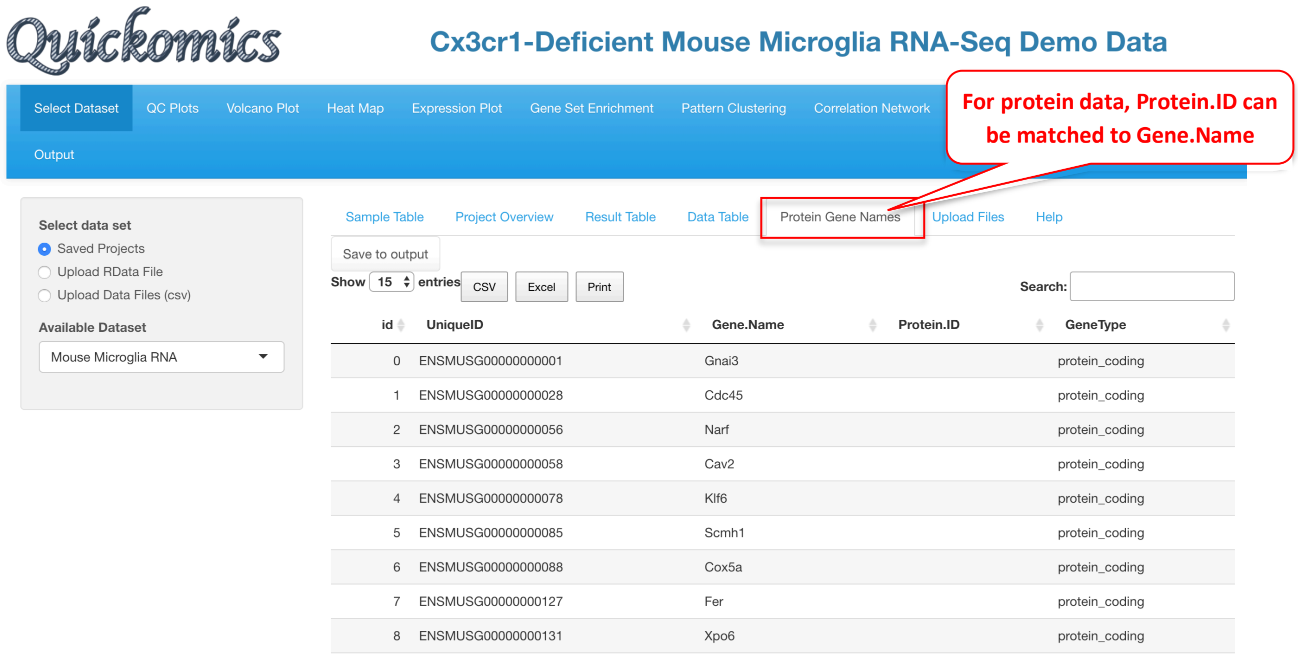

#### 2.8 Help

Finally, the "Help" feature summarizes what data is being displayed in each tab under Select Dataset. The Help tab is available for all sections of the subsequent analysis.

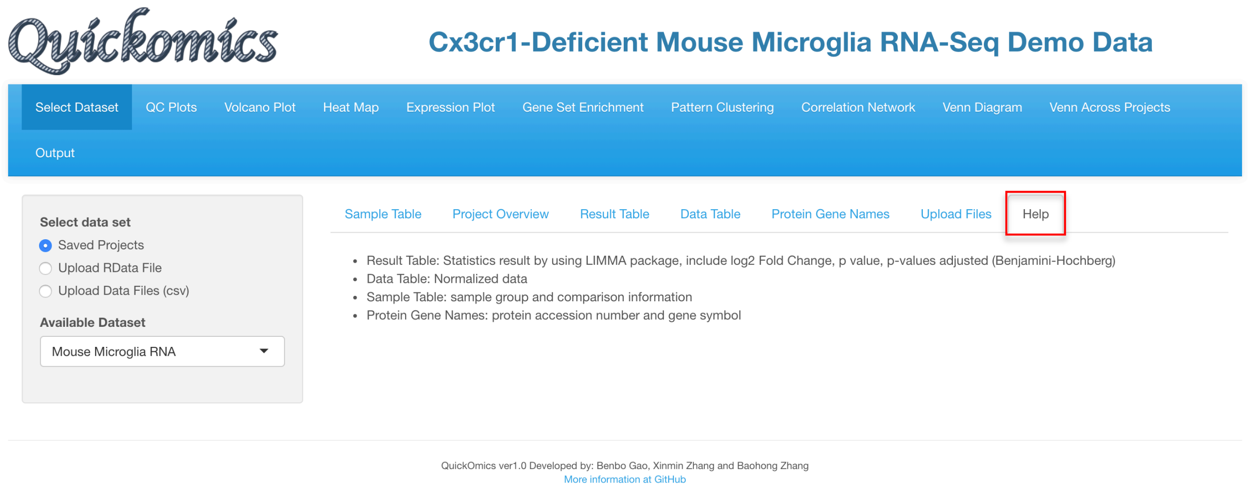

### 3 QC Plots Module

The “QC Plots” module performs basic QC analyses, as described in following paragraphs, on an uploaded dataset to evaluate integrity and quality of the data. The plots were generated using ggplot, plotly, ComplexHeatmap and pheatmap R packages.

#### 3.1 PCA Plot

The principle component analysis (PCA) plot displays, by default, the first 2 PC’s in the dataset and colors study groups with “none” selected in shape and size settings. This plot is highly customizable. Users can change the display by annotating the plot with different colors, shapes, size scheme, as well as label selected subsets data points.

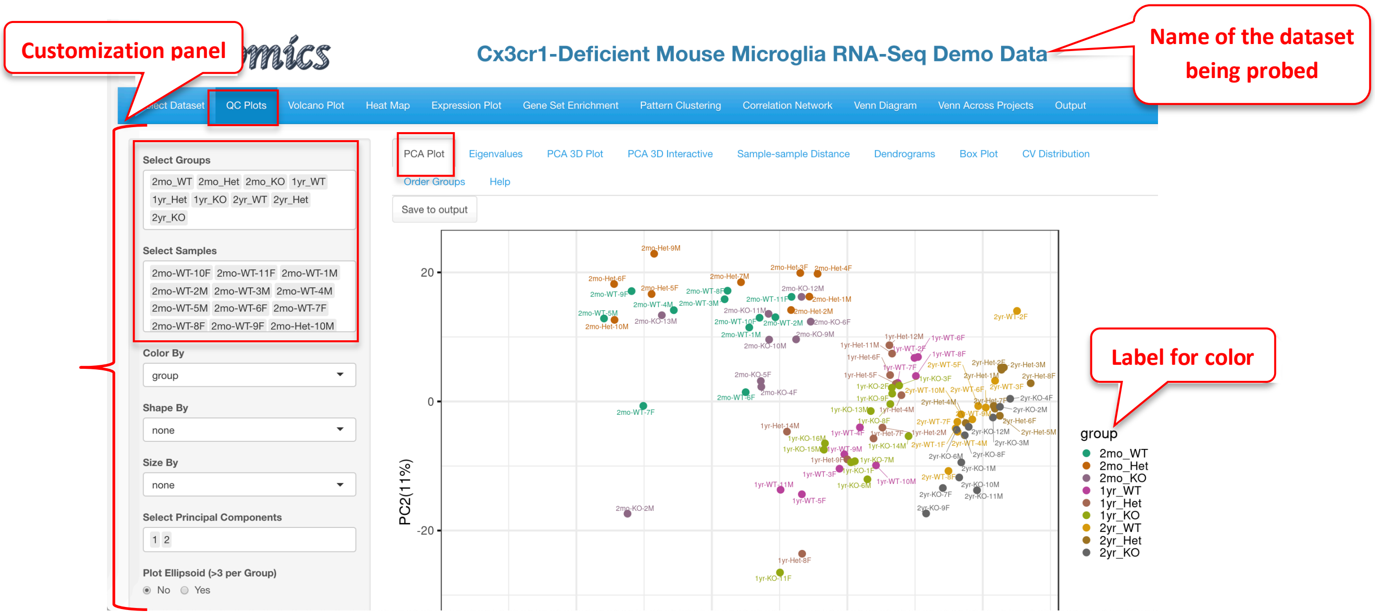

Additionally, the “Select Groups” and “Select Samples” feature (Green square) is present in all the QC Plots tabs. The User Is able to remove any particular group or sample from the plots.

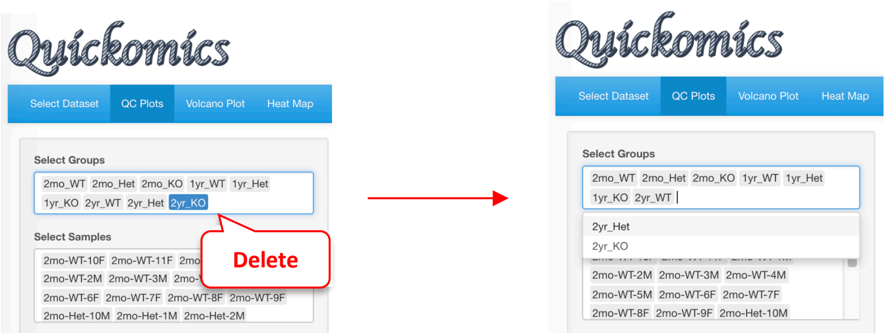

To illustrate the strengths of customization, we changed a few display elements as shown below, to visualize the influence of Age and Genotype on sample clustering:

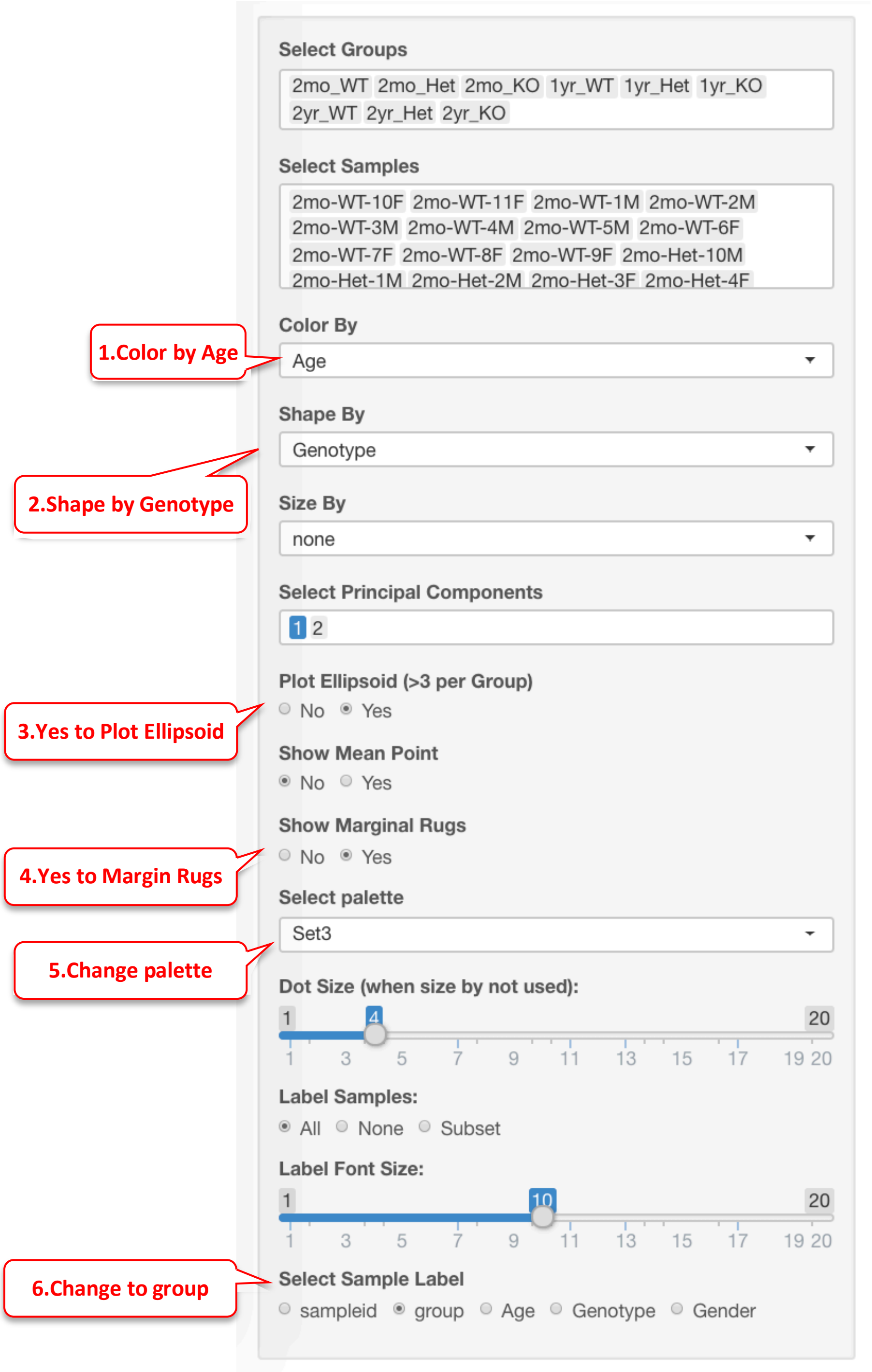

These changes results in a PCA plot that clearly showed that Age is the factor that drives the biggest separation, especially along PC1. There is some separation driven by the Genotype as well. The Ellipsoid feature helps group samples by the color attribute, while the Marginal Rug feature helps understand the sample distribution along the axis.

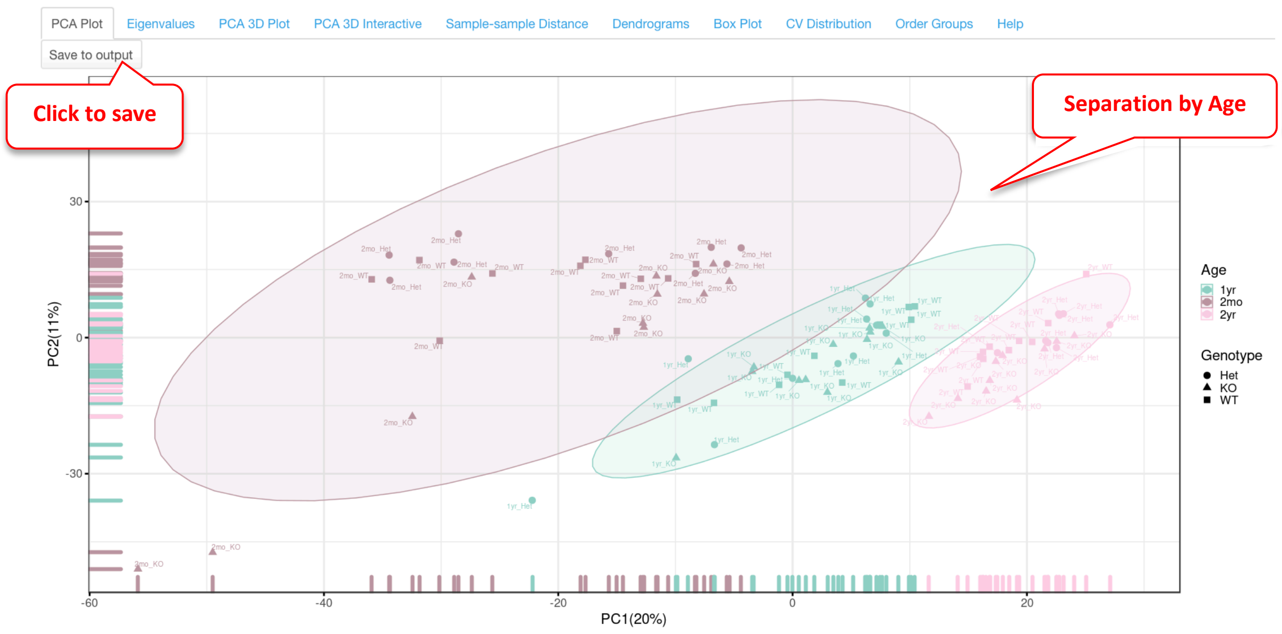

Furthermore, the plot can be saved in high resolution by clicking on “Save to Output”. Section 12 of this document describes the next steps of downloading and obtaining the saved plots.

#### 3.2 Eigenvalues

By default, the PCA plot displays PC1 and PC2 as described in Section 3.1. However, in many cases other PC's may be important in explaining additional variance in the dataset. This sub-tab generates a plot of the variances versus the first 10 PCs in the dataset, enabling Users to make educated decisions of which PCs to plot in 2D scatter plot. In this dataset PC1 and PC2 together explain around 30% of the variance.

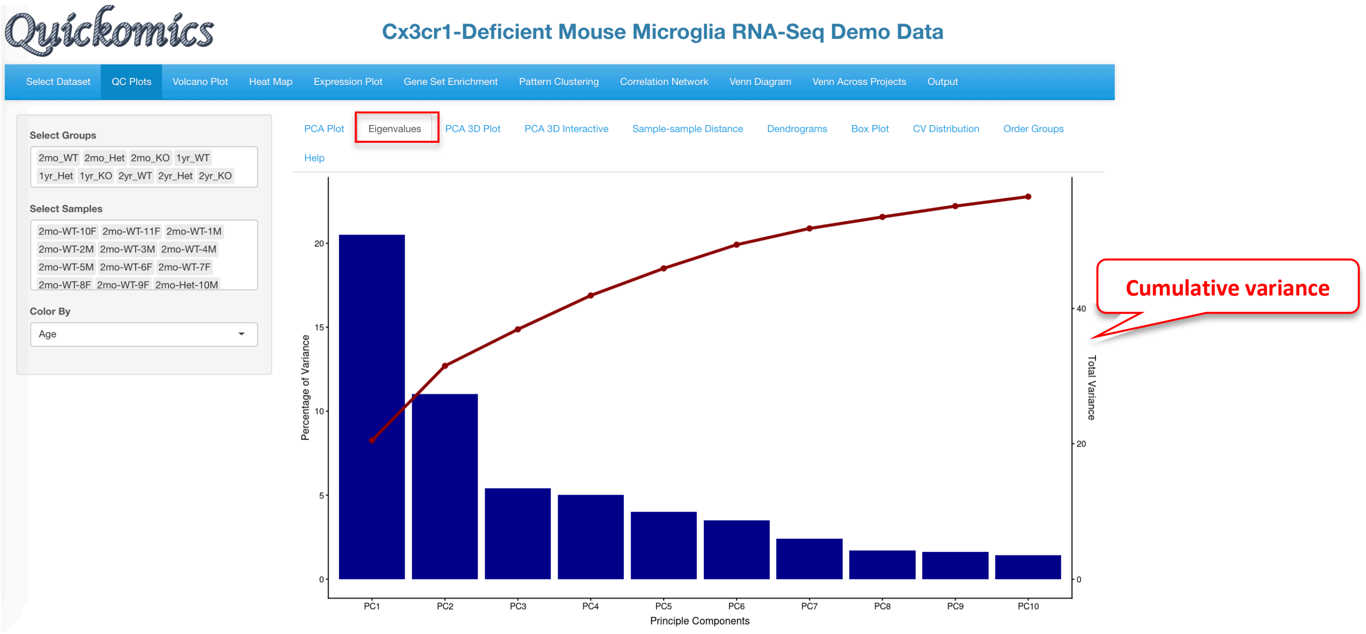

PC2 and PC3 were used to redo the PCA plot, but here the factors driving the separation of the samples is not as clear.

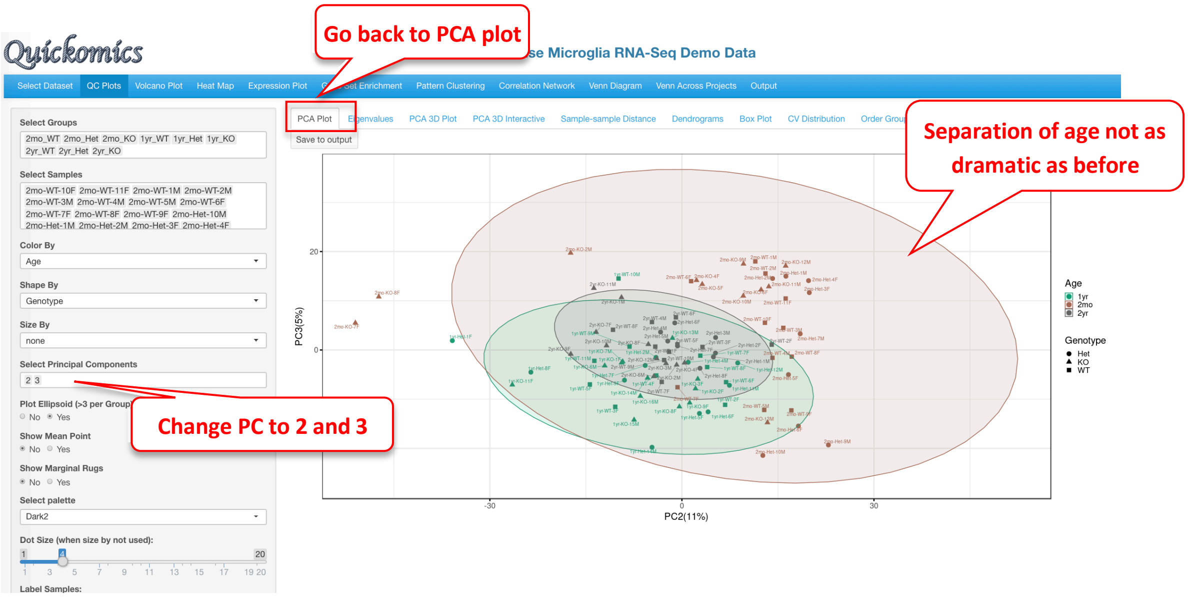

#### 3.3 PCA 3D Plot

This sub-tab contains a way to perform 3D PCA representation. The functionality is limited to doing so only on PC1, PC2 and PC3. Users can zoom in and out to focus on different areas of the plot or rotate the plot in different axes to find the best separation by age as shown below. There are 3 attributes that can be changed: color, ellipsoid inclusion, and labels of the samples.

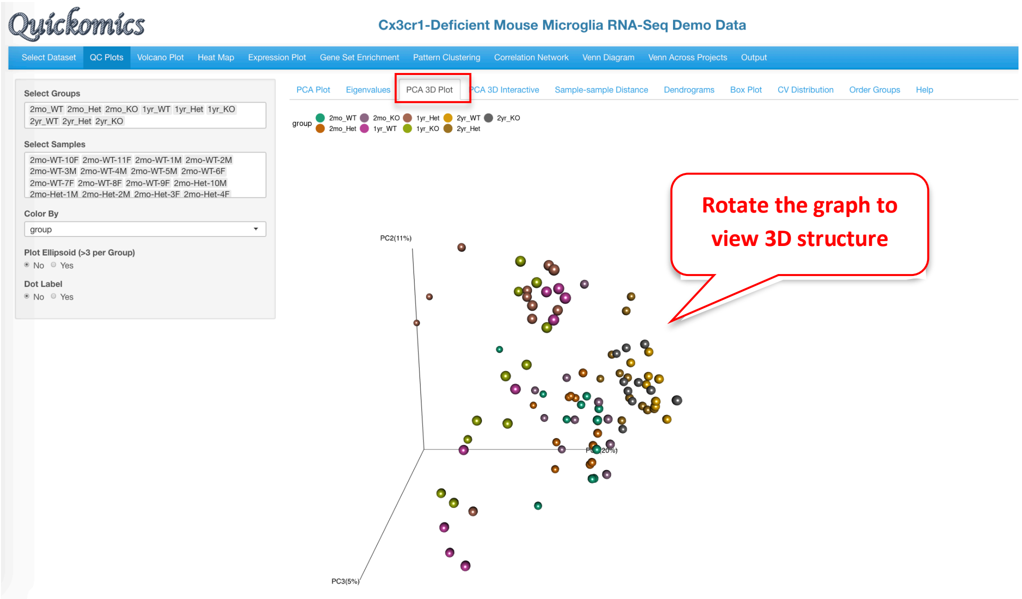

The "Color By" and "Plot Ellipsoid" attributes were changed in the following plot. This highlights the clear separation of samples by Age and not Genotype, contrary to the original biological hypothesis.

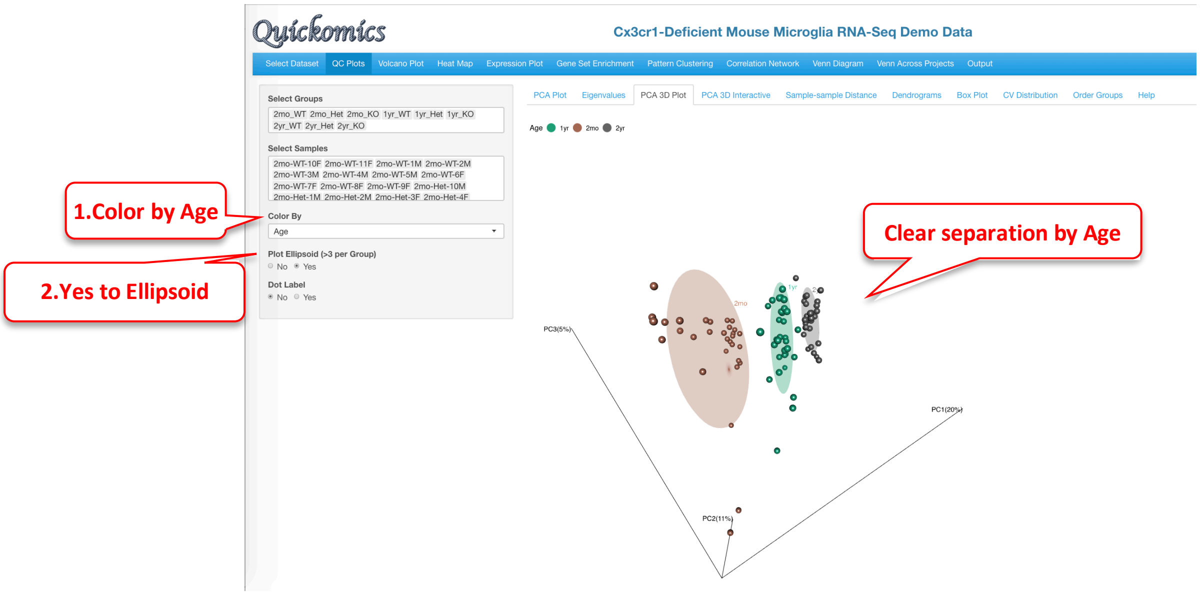

#### 3.4 PCA 3D Interactive

This tab is similar to the previous one (3.2 PCA 3D Plot), but here users have the additional capability to hover over the samples on the plot to identify more details, see the pointer below. This was implemented through the plotly package in R. Users can change 2 attributes here, the color and shape.

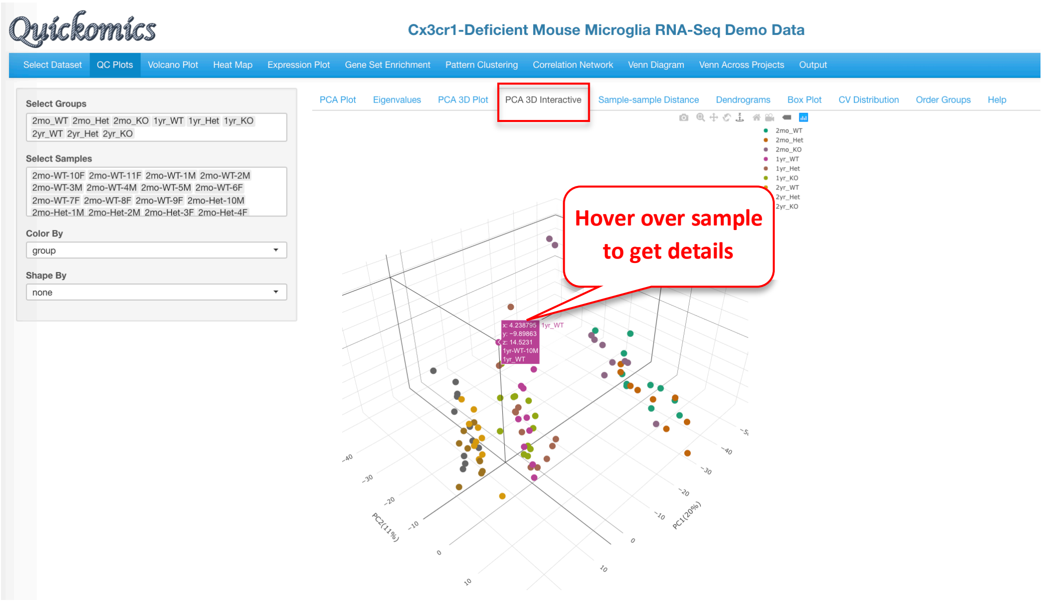

In the plot below, the “Color By” and “Shape By” attributes have been changed to highlight the drivers of separation.

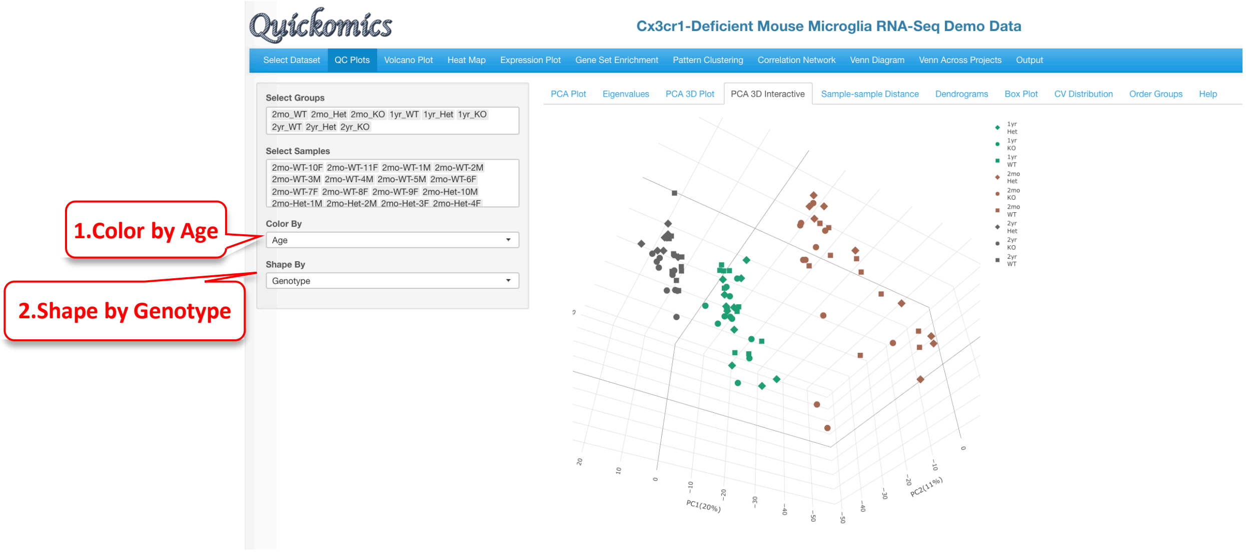

#### 3.5 Sample-sample Distance

This tab helps identify pair-wise similarity between all samples. A distance matrix is generated for every sample pair and is plotted as a heatmap. The rows and columns are clustered based on similarity.

It is clear from this plot that samples of the same Age have the smaller distances, that is, they have closer gene expression patterns. Here again, Users have the option to save to output.

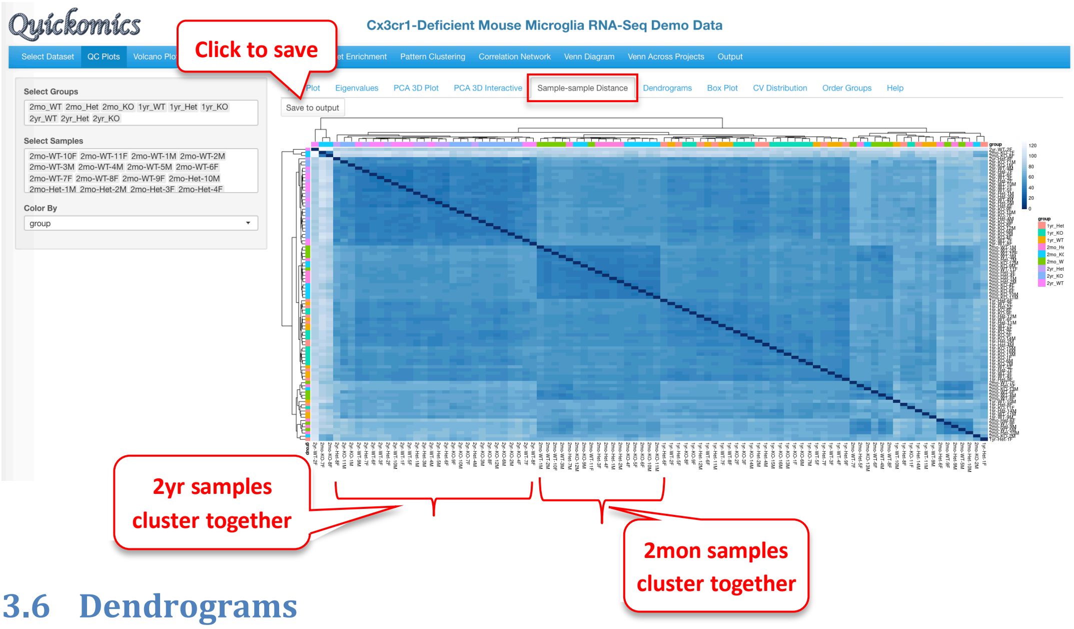

#### 3.6 Dendrograms

Like the previous plot (3.5 Sample-sample Distance), the Dendrogram plot helps visualize hierarchical clustering relationships between samples. The Default plot is Circular and cut into four parts. Users have the option to visualize in two other ways (tree or horizontal) and cut the plot in multiple regions. Here again, Users can save the plot to output.

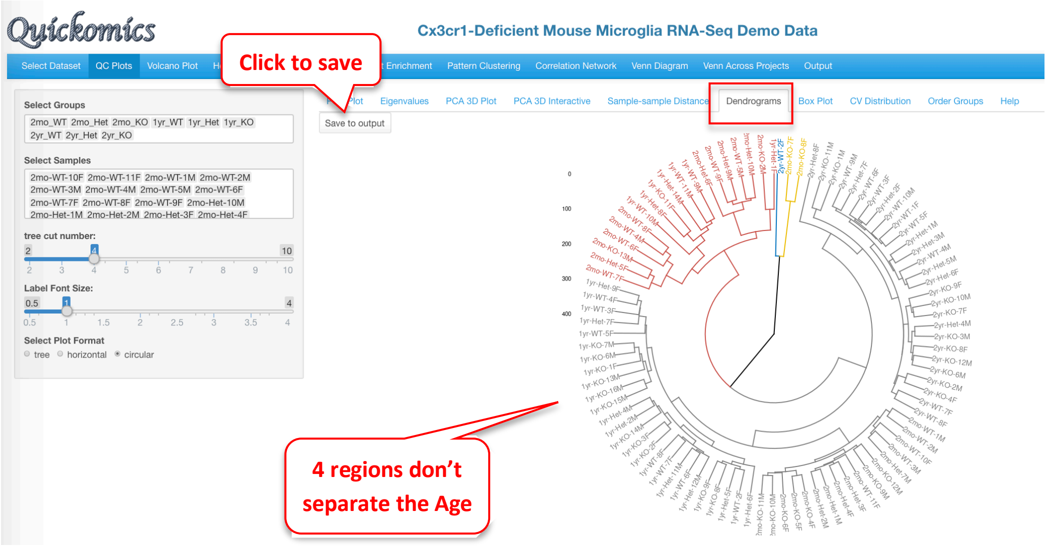

In the plot below, we have selected to visualize as a tree and cut into eight parts. This shows the relationship between the samples to help understand the hierarchal clustering. Overall, the samples are arranged by age, but there are some outliers that appear. It is also interesting how the 2mo samples cluster with the 1yr samples.

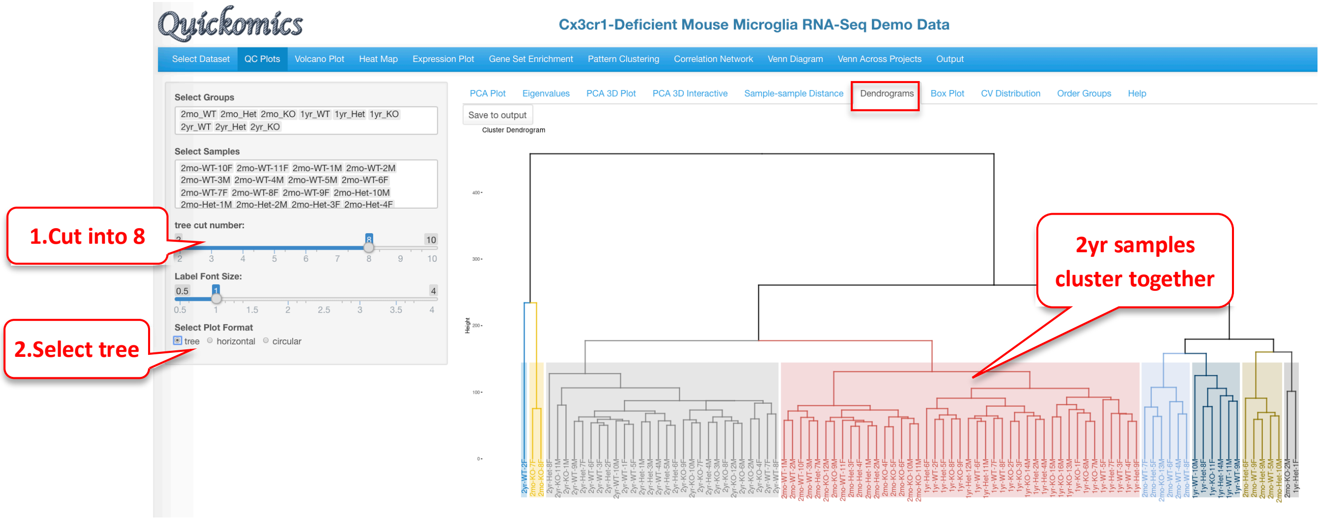

#### 3.7 Box Plot

The Boxplot is a visualization to understand the distribution of the normalized expression levels in all samples. This identifies the minimum, first quartile, median, third quartile, and maximum values in the dataset. In this demo dataset, most of the samples have the same range of expression, indicating that there are no outliers.

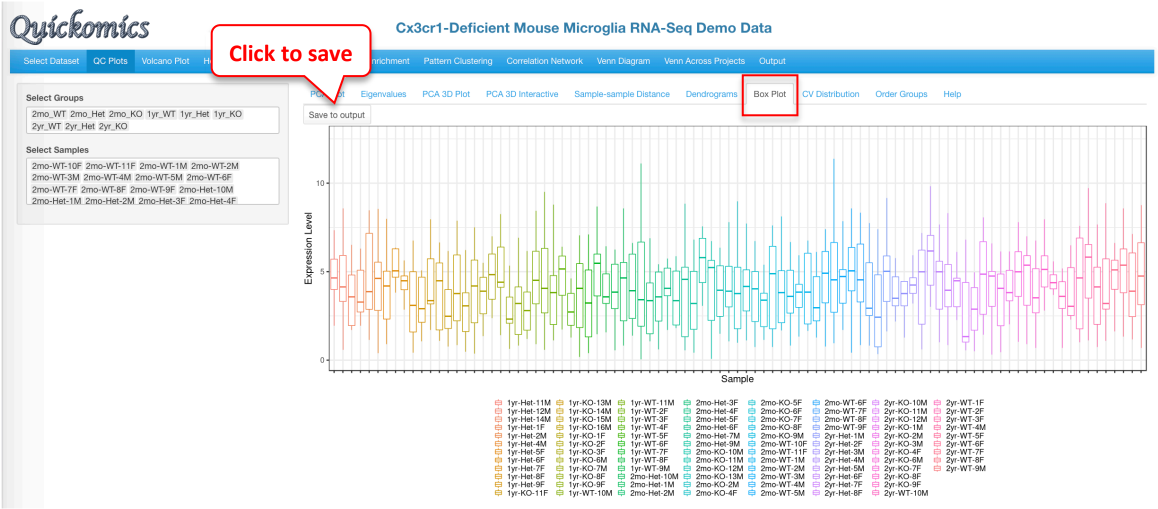

#### 3.8 CV Distribution

This plot shows the histogram of CVs (coefficient of variation) and a dotted line for each group indicates the median CV. In this dataset, the 2mon_KO samples have the highest CV while the 2yr_Het samples have the lowest variability, which corresponds to what is visually seen on the PCA plot for these samples in Section 3.1.

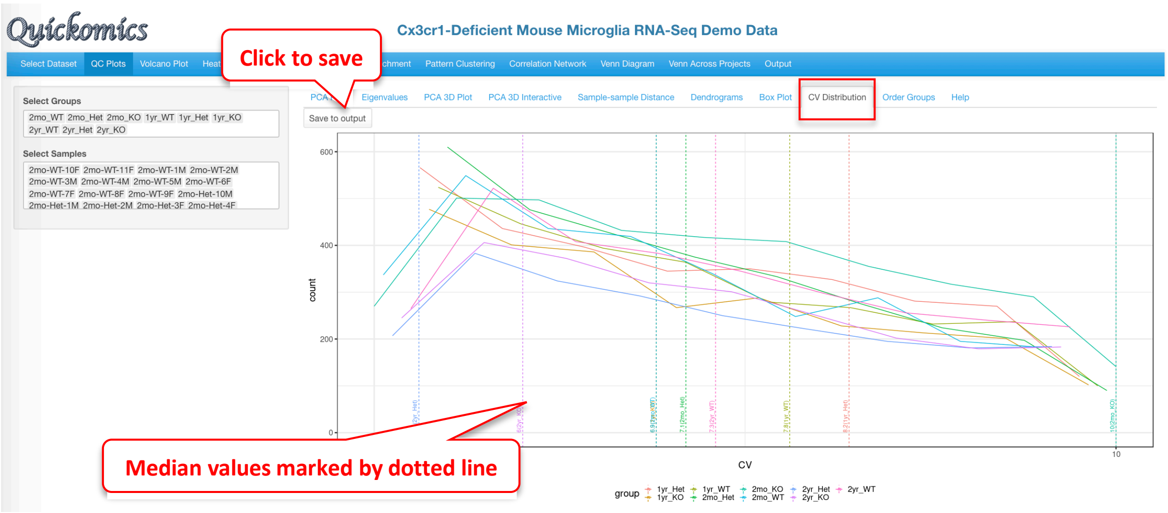

#### 3.9 Order Groups

In this sub-tab, Users can select and/or re-order the groups in the dataset. The order will be stored in the current session, and labels will be displayed in this preferred order for all plots. Users can either drag and drop a group to reorder or use the select panel to delete and add. In this example the KO genotype was of highest interest and so we have changed the order of the group so that the KO samples are before Het and WT.

| Drag and drop | Delete and Add |
|---|---|
| 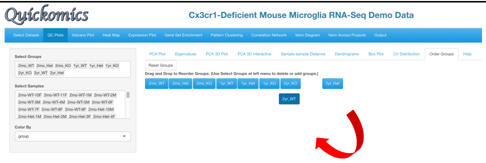 | 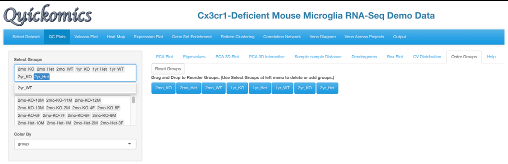 |

The changed groups are now visible. To revert back to the default, Users have the option to click "Reset Groups".

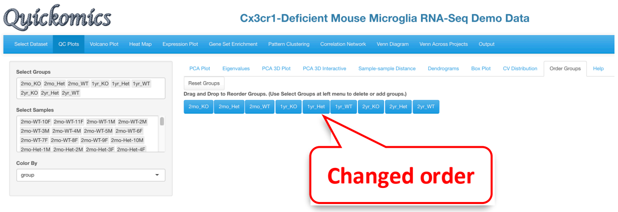

As noted above, that the new order of groups will dynamically change all other related visualizations. As an example, a 2D PCA plot reflecting a newly changed group order is shown below.

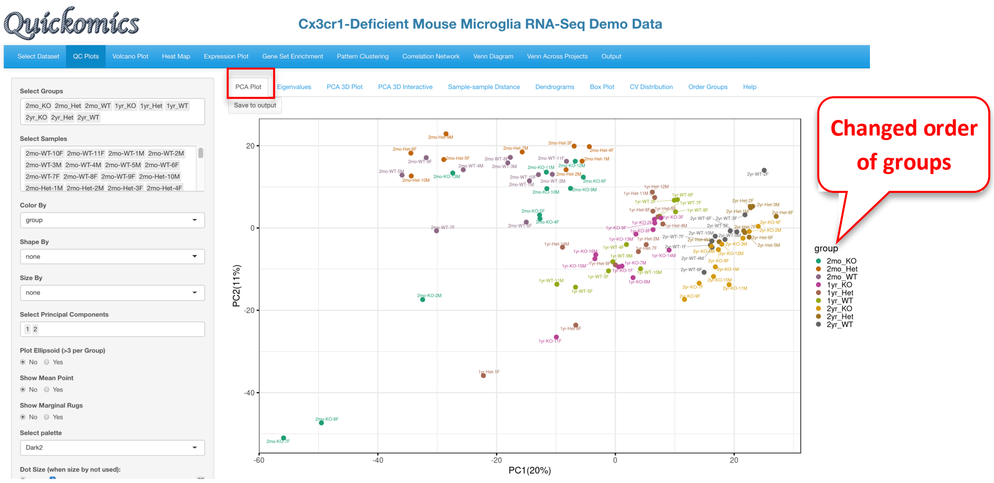

#### 3.10 Help

The Help section describes each tab of the QC Plot and what is being visualized. For further help please visit the GitHub containing the source code for this tool (https://github.com/interactivereport/Quickomics) or click on the link at the bottom of the page.

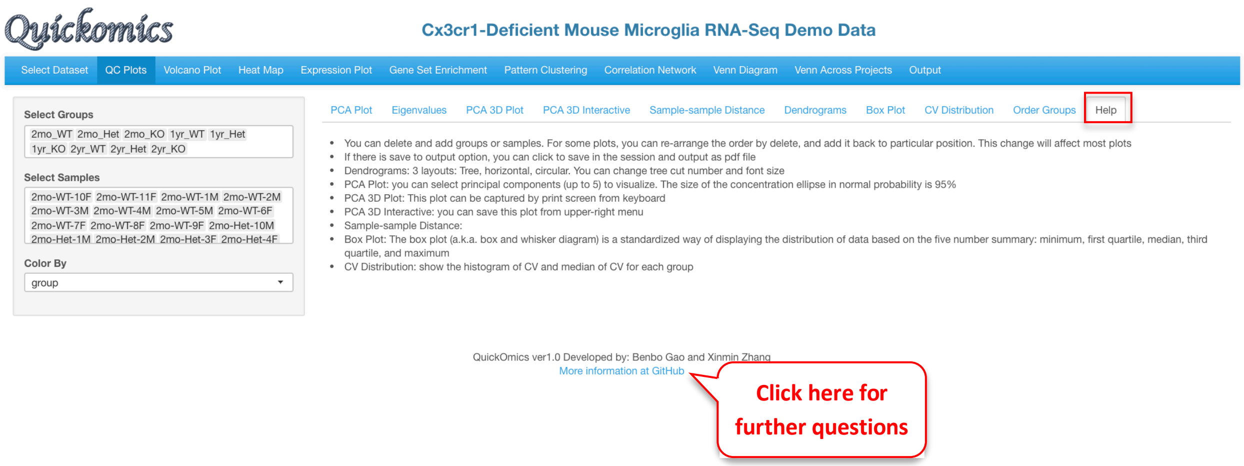

### 4 Volcano Plot Module

This section is designed to explore the differentially expressed genes (DEGs) and differentially expressed proteins (DEPs) in the dataset. The comparisons are defined by the User when they input the dataset. Please refer to our GitHub for more details on uploading datasets.

The “Volcano Plot (Static)” described in Section 4.1 and “Volcano Plot (Interactive)” described in Section 4.2 are ways to visualize each pair-wise comparison one by one. “DEGs/DEPs in Two Comparisons” described in Section 4.3 is a novel way to visualize the similarities and differences in two comparisons. Finally, “Data Table” described in Section 4.4 displays a searchable table containing all genes and their fold changes and P values in each comparison.

#### 4.1 Volcano Plot (Static)

This plot reveals the differentially expressed genes provided in the input data file. By default, the Fold change Cutoff is set to 1.2 and the P value Cutoff is set to 0.01. Users have the option to change any of these parameters. For more advanced display options, they can set the “Show More Options” to “Yes” to customize other settings.

The red dots are indicative of all genes that pass the Fold Change and P value cutoff, while the green dots are indicative of genes that only pass the P value cutoff. Fifty random genes are labelled for both the up-regulated and down-regulated genes by default.

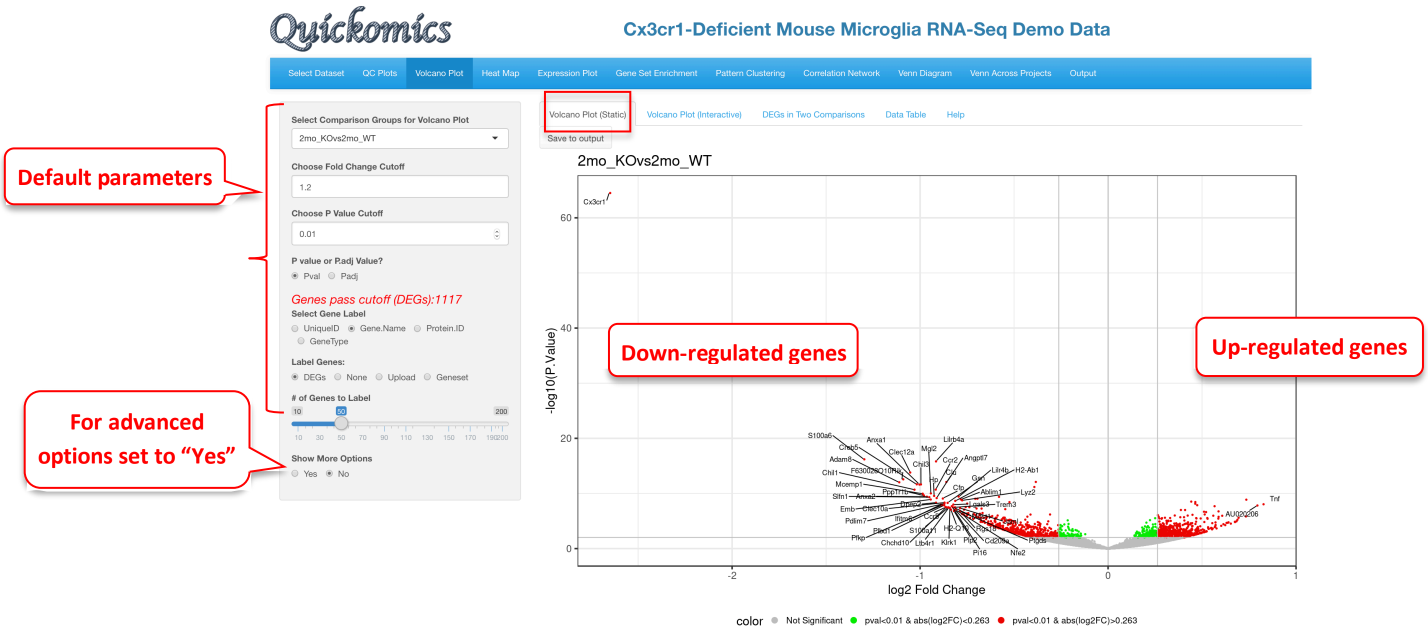

For demonstration purposes, we have changed the following attributes to highlight a few genes in this dataset. We have selected the 2mon_KOvs2mon_WT comparison, where Gyoneva et al., 2019 report immune response genes as the top differentially expressed category. To highlight these genes, in the example below, we select an “Immune Response” dataset from the Molecular Signatures Database (MSigDB) to highlight the genes by adding labels to the volcano plot. As MSigDB contains only human gene lists and the demo dataset is from mouse, Quickomics does a quick conversion of the names of

Human genes to Mouse genes by changing the letters after the first one to lowercase. Additionally, in this example we removed some of the genes from the list to reduce size. This visualization was not available in the original publication, but clearly supports the finding that the KO genotypes altered the expression of immune-related genes.

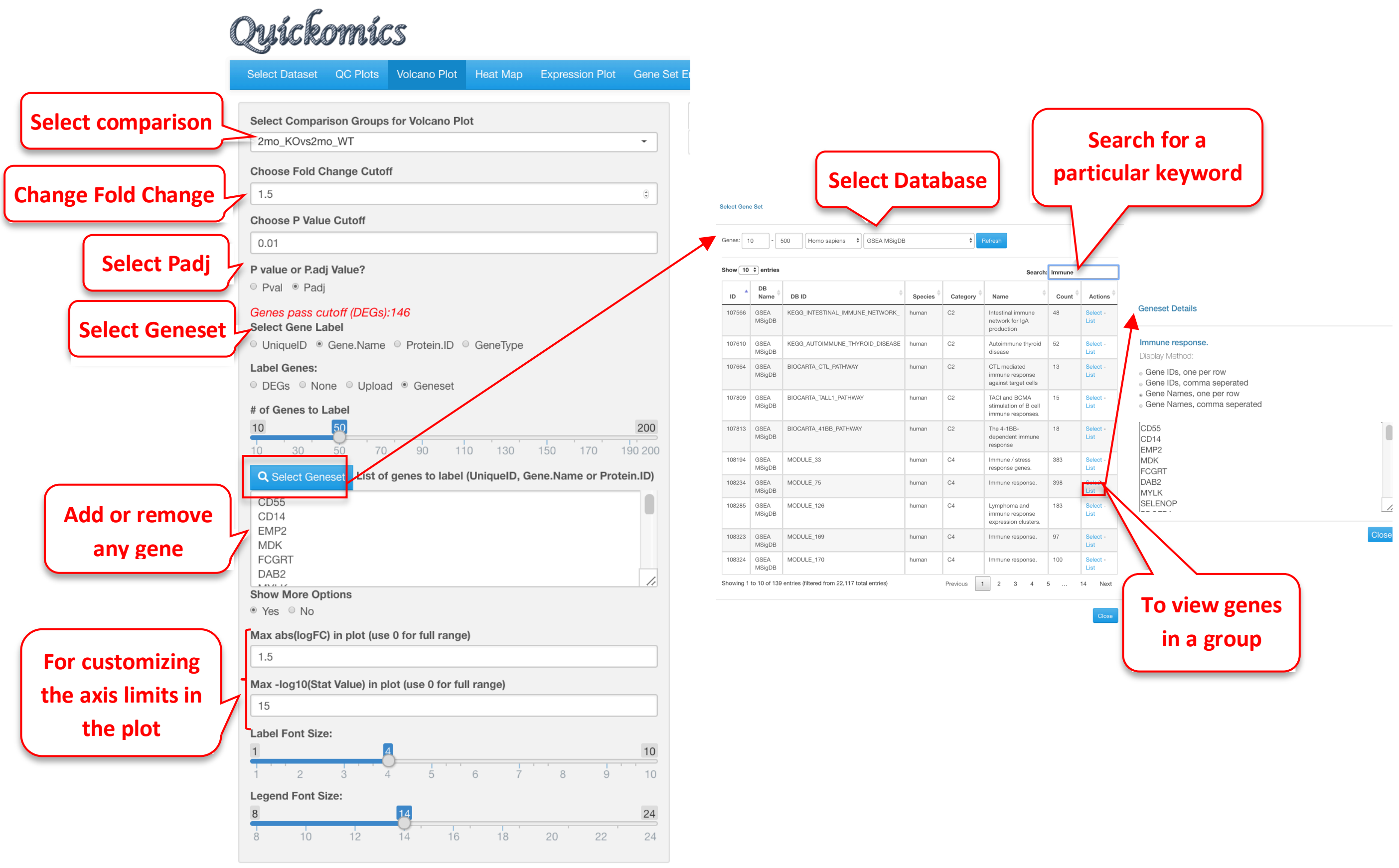

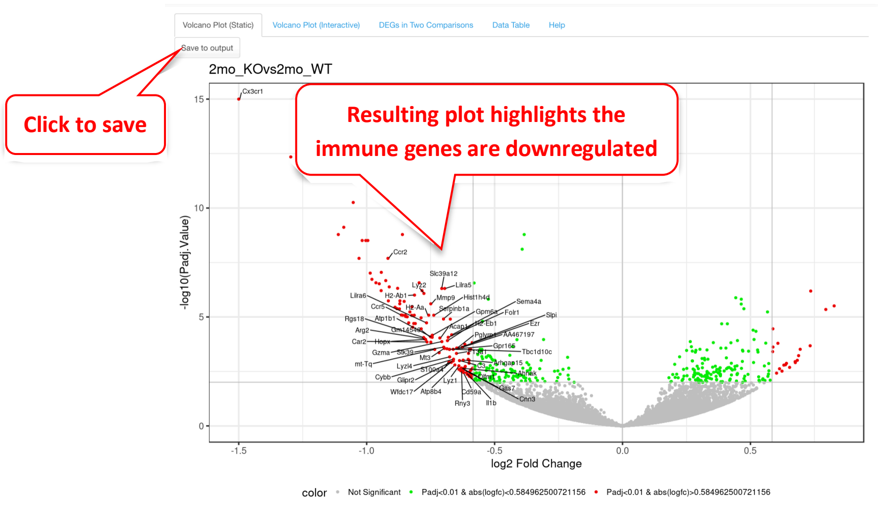

#### 4.2 Volcano Plot (Interactive)

This next sub-tab is a very similar visualization as the previous one (4.1 Volcano Plot (Static)) with an additional capability where users can hover over a particular dot/gene on the plot to see more details. This plot has a few attributes that can be changed, and in the example below we have used the same cutoffs and values as in section 4.1.

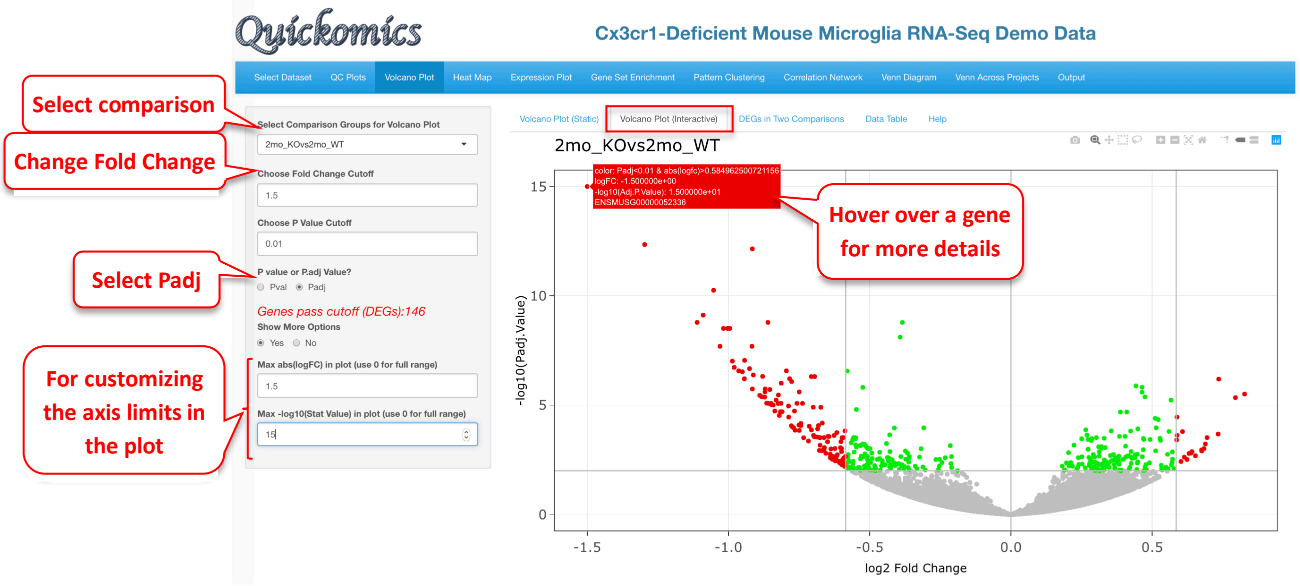

#### 4.3 DEGs and DEPs in Two Comparisons

This next tab helps with identifying DEGs/DEPs that follow a similar trend in two comparisons. Gyoneva et al., 2019, the source of this demo dataset, report that the genes differentially expressed in Cx3cr1 KO microglia in the 2mon KO are quite different from WT, while the differences from WT are smaller for the 1yr KO and 2yr KO, and the two older ages are similar to each other. We have plotted out the following comparisons that illustrate this conclusion.

2mo_KOvs2mo_WT vs 1yr_KOvs1yr_WT has a low correlation score. There are many genes that are colored yellow, meaning those genes were significant in 2mo_KOvs2mo_WT (Y axis) but not in 1yr_KOvs1yr_WT (X axis).

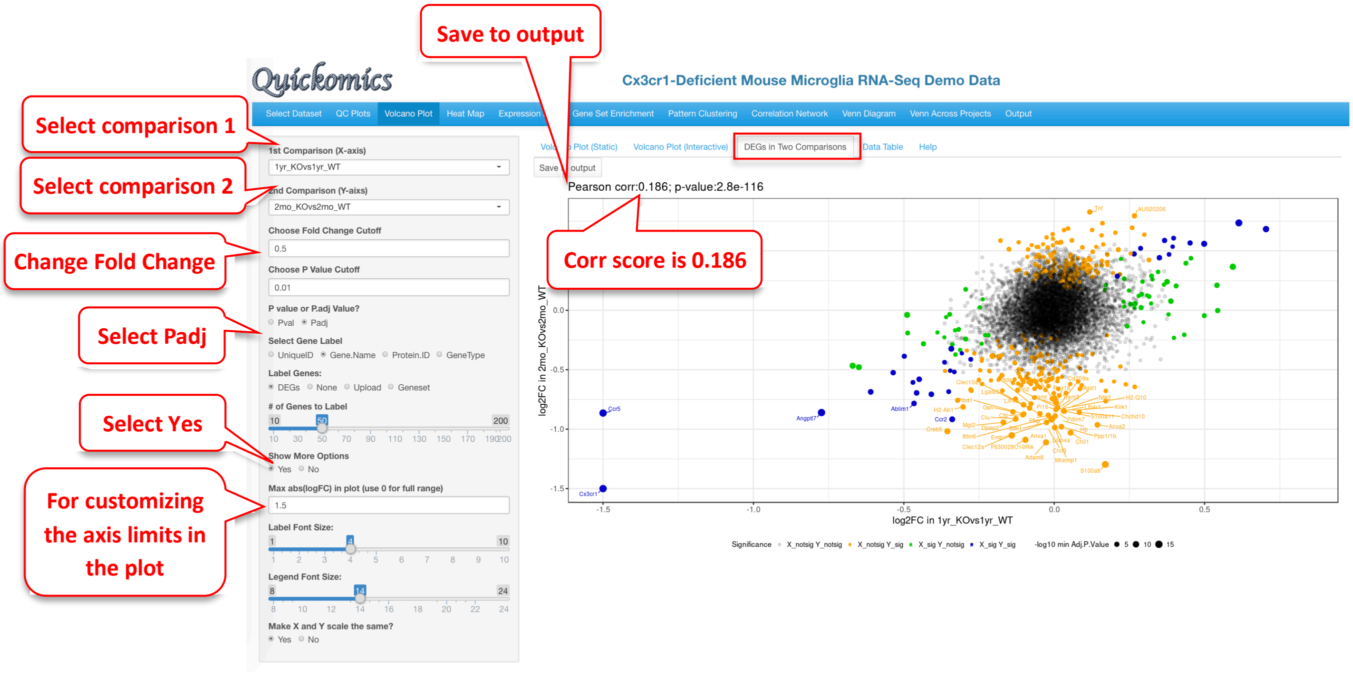

Similarly, we changed the X axis to 2yr_KOvs2yr_WT to compare to 2mo_KOvs2mo_WT, and it shows a very similar plot with a low correlation score of 0.178

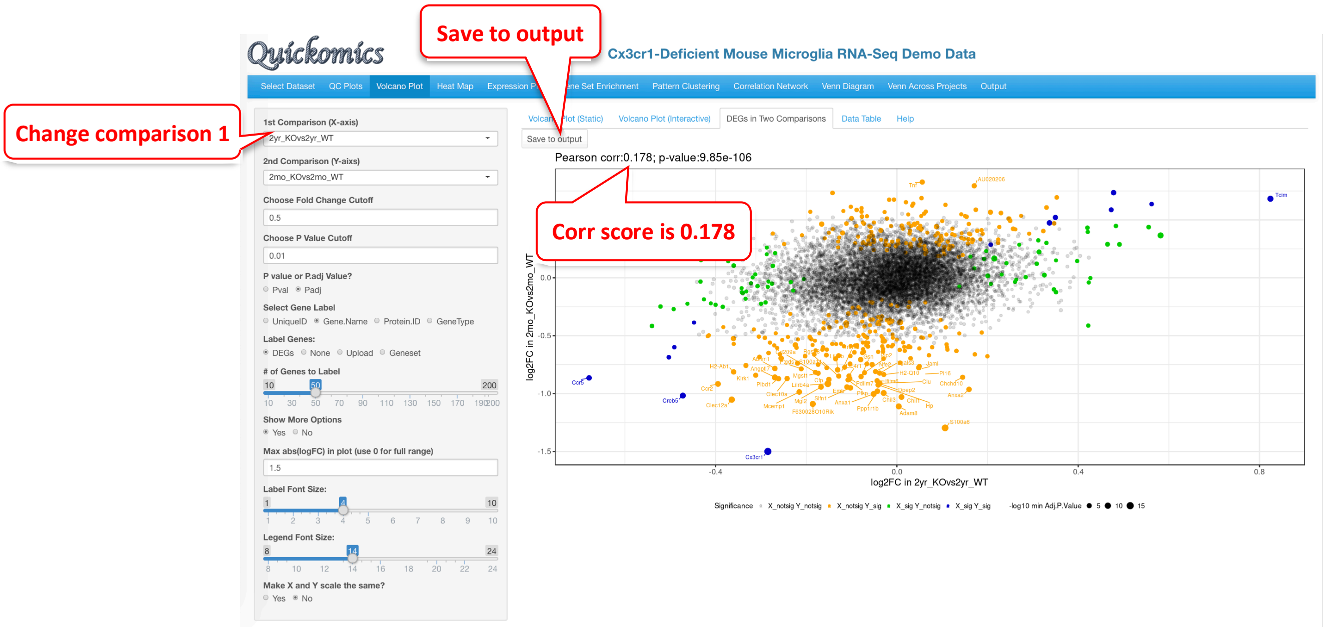

Next, we changed the Y axis comparison to 1yr_KOvs1yr_WT comparing it to 2yr_KOvs2yr_WT on the X axis. This plot has a higher correlation score of 0.371 than the previous one, suggesting that these two comparisons are more similar. These comparisons represent additional insights supported by statistical analyses that were gained from the data and were not available in the original manuscript by Gyoneva et al.

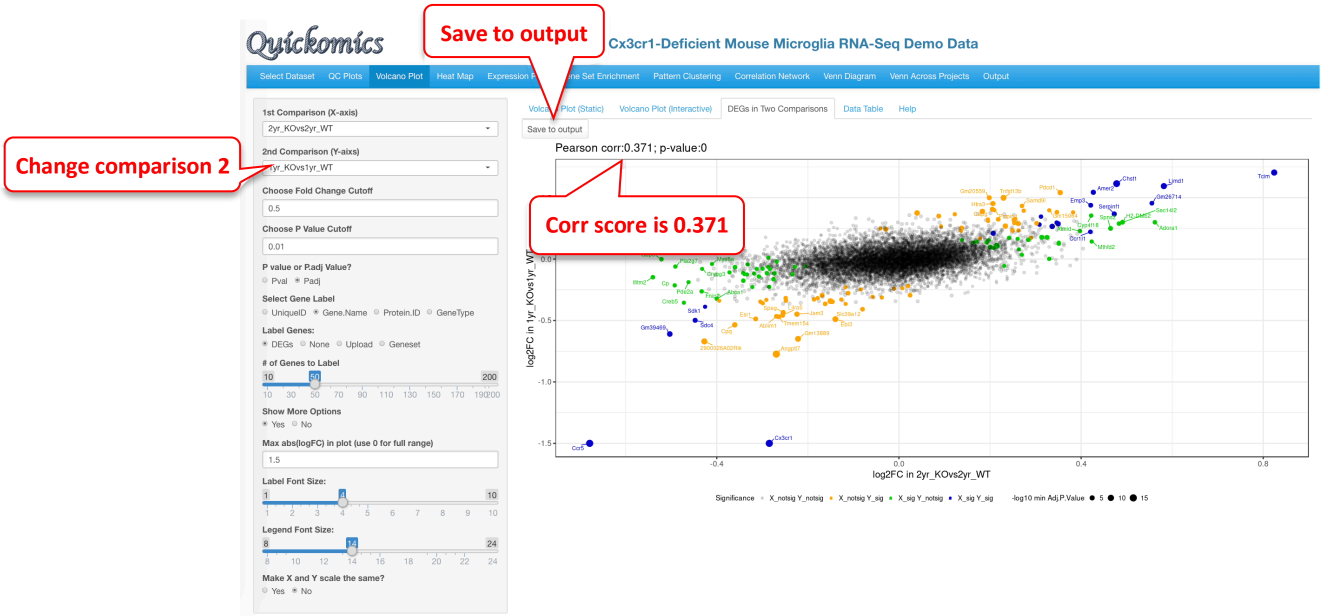

#### 4.4 Data Table

This tab lets the users view the DEGs/DEPs in a tabular format with a searchable feature. Users also have the ability to download the gene list as a CSV.

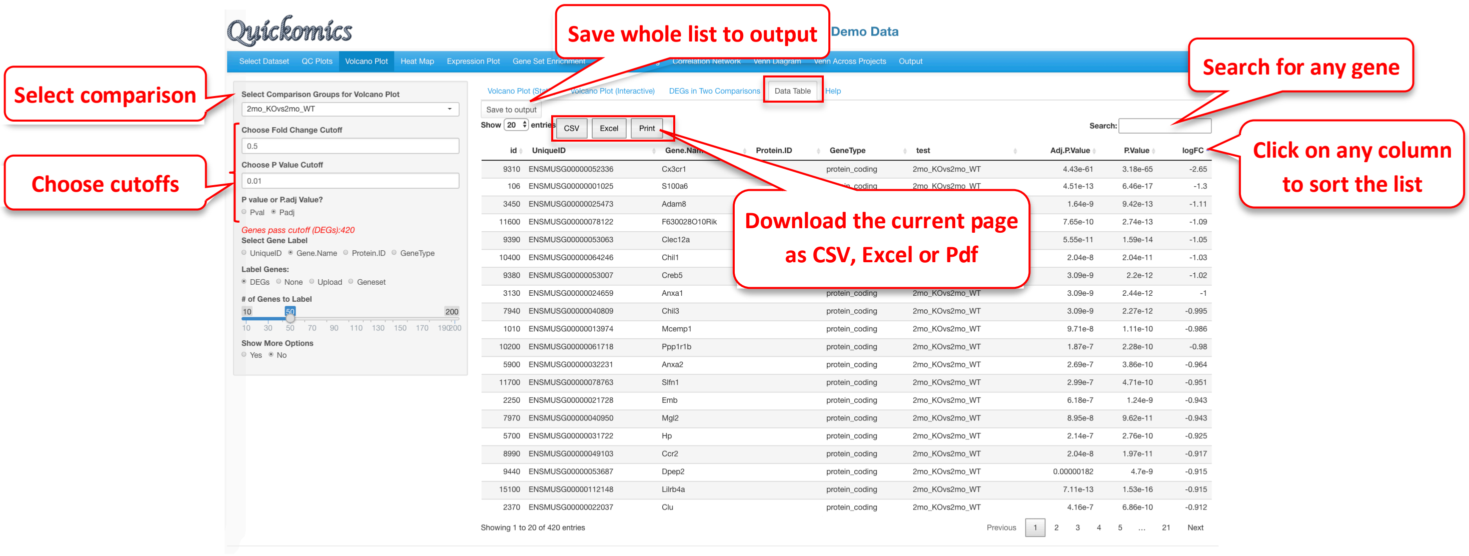

### 5 Heatmap Module

This module generates highly customized publication-quality heatmaps with the ComplexHeatmap package in R. Users can enter automated list of differentially expressed genes or enter a custom list of genes to highlight on the heatmap. This module differs from other modules in that each time Users change the attributes; they need to click on the “Plot/Refresh” button to generate the plot. This is to avoid changing the output before all the attribute are added.

#### 5.1 Static Heatmap Layout 1

The “Static Heatmap Layout 1” uses the ComplexHeatmap package (Gu *et al.*, 2016) for generating the plot. For demo purposes, we have selected options that highlight some capabilities to generate informative heatmap. For gene selection the default option is the selection of 100 random genes which can also be changed to variable genes. Here we selected DEGs/DEPs from the 2yr_WTvs2mon_WT comparison. Users also have the option to change the cutoff criteria for filtering the genes. They can also add/remove annotation categories for the samples on the columns.

In this heatmap, the top up-regulated genes in 2mon_WT samples are seen as high in 2mon samples but low in 2yr and 1yr samples, while the top up-regulated genes in the 2yr_WT samples are high in the 2yr and 1yr samples but low in the 2mon samples. This is another example that age has a major effect on gene expression in microglia, and the 2mon microglia have a different gene expression profile compared to 1yr and 2yr.

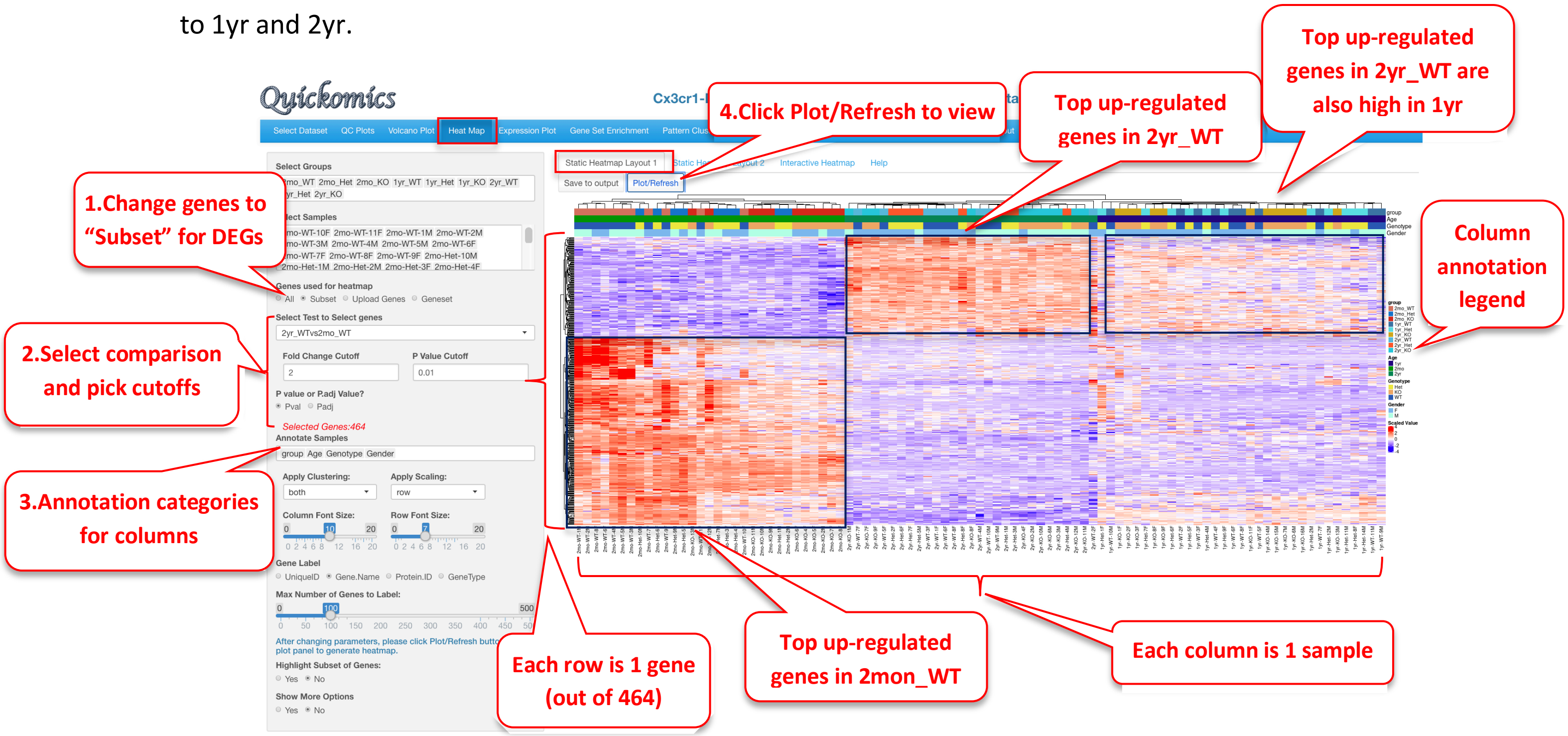

In this next example, we customize another Heatmap to highlight immune related genes that were called out in Gyoneva et al., 2019. This is done by uploading a csv file consisting of the gene names with pathway information and the color for display. An example of this file is on our GitHub (Quickomics/demo_files/fig5_all_genes.csv)

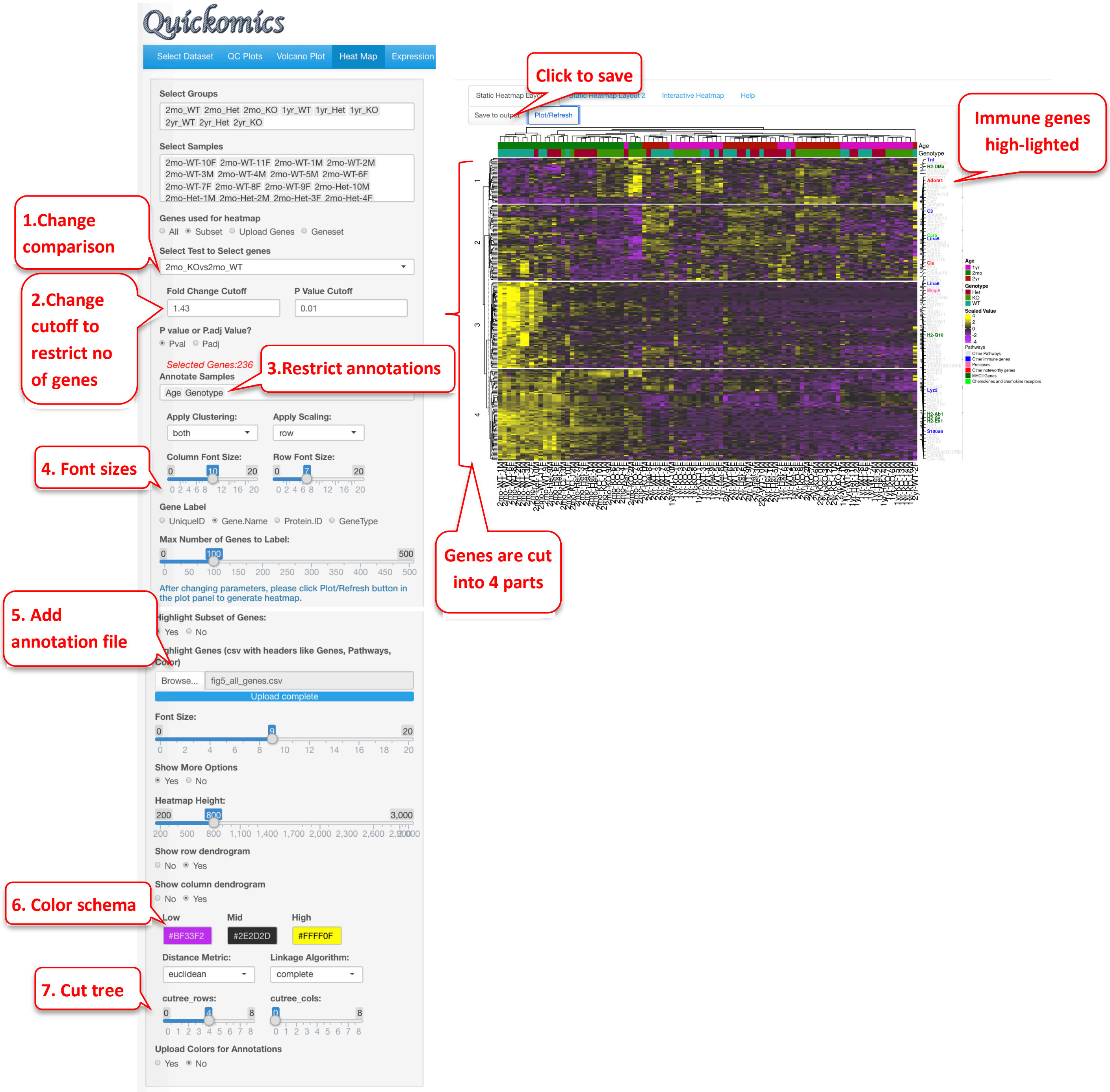

#### 5.2 Static Heatmap Layout 2

The "Static Heatmap Layout 2" uses the heatmap.2 function from the gplots package for generating the plot. Here is a demo of the same set of genes with dendrograms for both rows and columns.

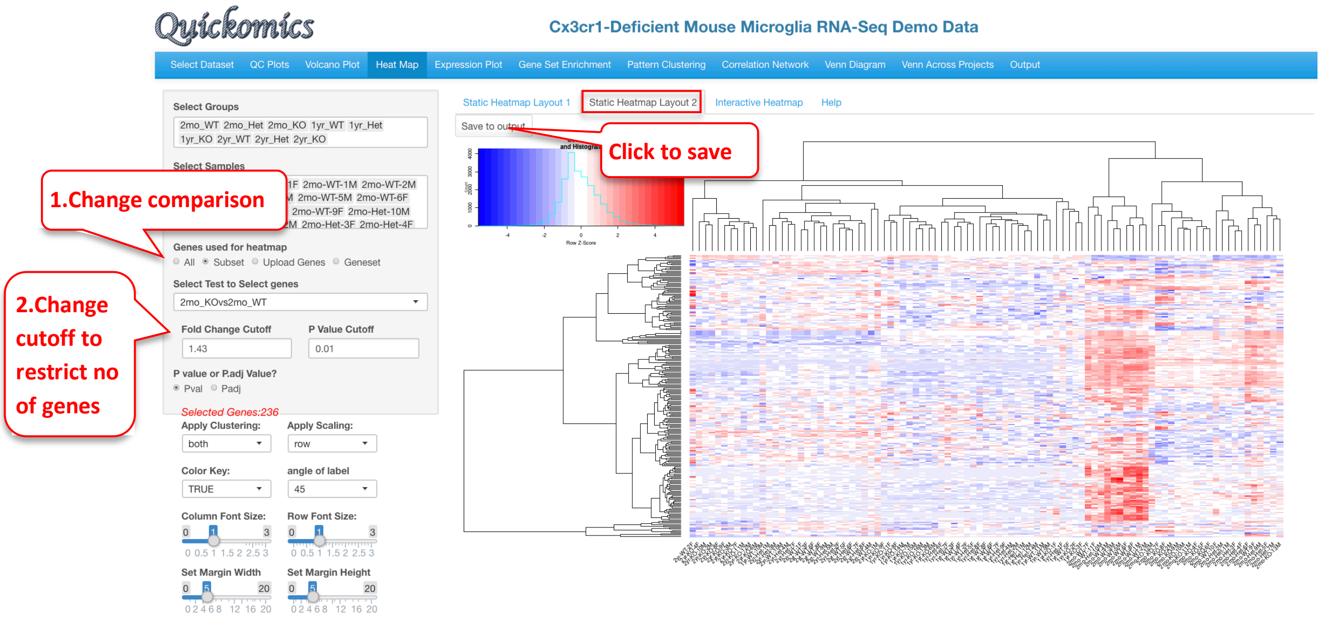
Quickomics
Cx3cr1-Deficient Mouse Microglia RNA-Seq Demo Data
Select Dataset
QC Plots
Volcano Plot
Heat Map
Expression Plot
Gene Set Enrichment
Pattern Clustering
Correlation Network
Venn Diagram
Venn Across Projects
Output
Select Groups
2mo_WT 2mo_Het 2mo_KO 1yr_WT 1yr_Het 1yr_KO 2yr_WT 2yr_Het 2yr_KO
Select Samples
1.Change comparison
Genes used for heatmap
All Subset Upload Genes Geneset
Select Test to Select genes
2mo_KOvs2mo_WT
2.Change cutoff to restrict no of genes
Fold Change Cutoff
1.43
P Value Cutoff
0.01
P value or P.adj Value?
Pval Padj
Selected Genes:236
Apply Clustering:
both
Apply Scaling:
row
Color Key:
TRUE
angle of label
45
Column Font Size:
Row Font Size:
Set Margin Width
Set Margin Height
Static Heatmap Layout 1
Static Heatmap Layout 2
Interactive Heatmap
Help
Save to output
Click to save
Row Z-Score

### 6 Expression Plot Module

This functional module allows Users to browse through a subset of genes/proteins. It also offers an option to plot expression of individually selected genes. Four different displays: Box Plot, Bar Plot, Violin Plot and Line Plot are made available to fit the Users' preference.

#### 6.1 Browsing

This tab lets the Users plot out the expression of the DEGs/DEPs identified in the different comparisons. In this example below, we demonstrate how to display a few top genes as Violin Plots.

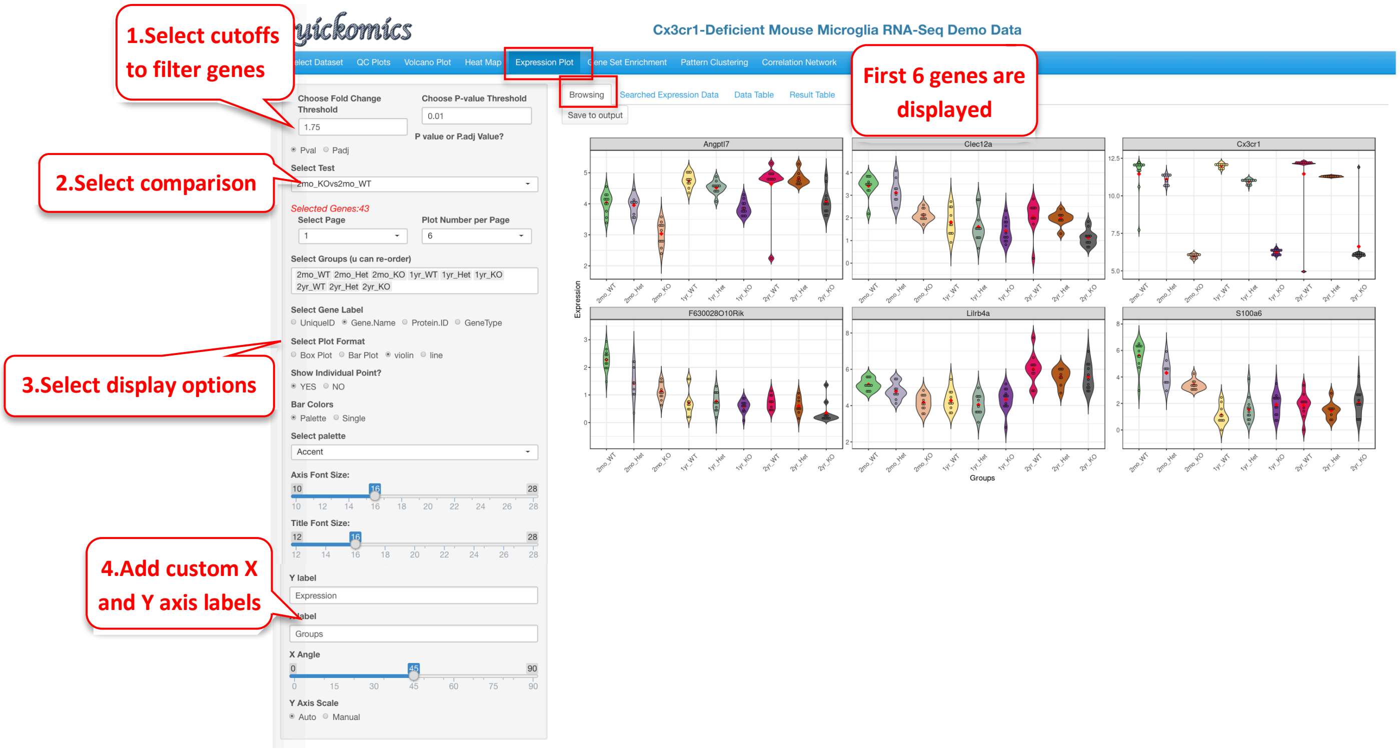

#### 6.2 Searched Expression Data

Another way to plot genes/proteins is using this functionality. Users can either enter a custom list of genes/proteins or select a list from online databases like KEGG, MSigDB etc. The selected genes/proteins can be plotted on a single plot or in multiple plots. Here is an example of plotting genes involved in MHCII antigen processing.

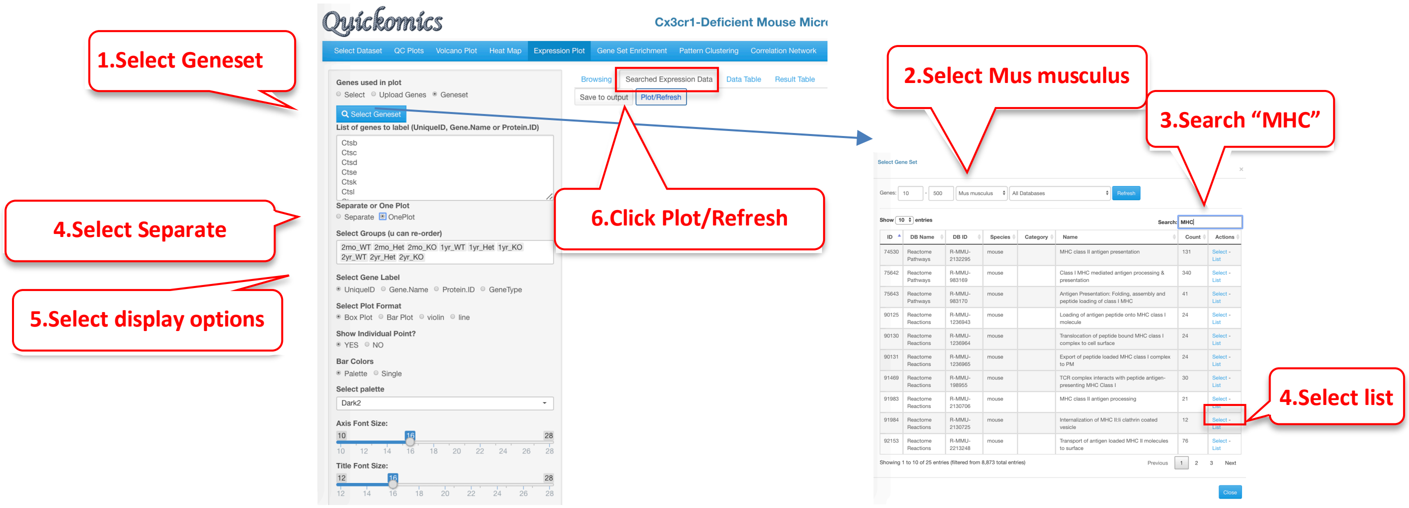

The resulting plot shows the expression of a subset of genes from the MHCII antigen presentation family of genes. Many of them appear altered by gene KO in 2mon microglia, but also in all genotypes in 2yr microglia, as indicated in the publication.

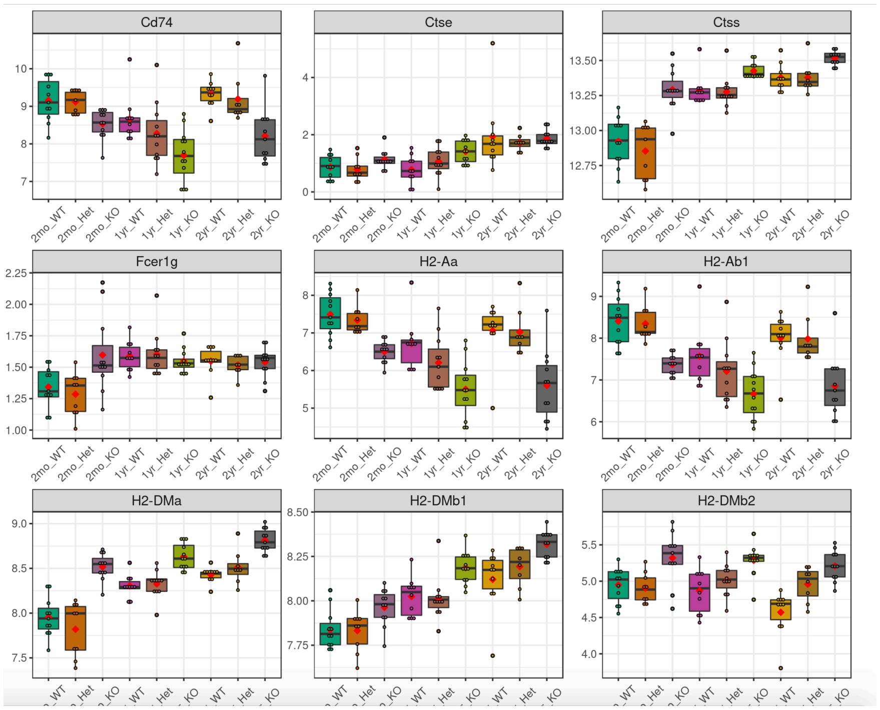

#### 6.3 Data Table

This sub-tab lists the normalized data of the genes/proteins selected in “Searched Expression Data” (Section 6.2) as a searchable table. For a demo of this feature, we select a proteome dataset from published paper (Connor-Robson *et al.*, 2019).

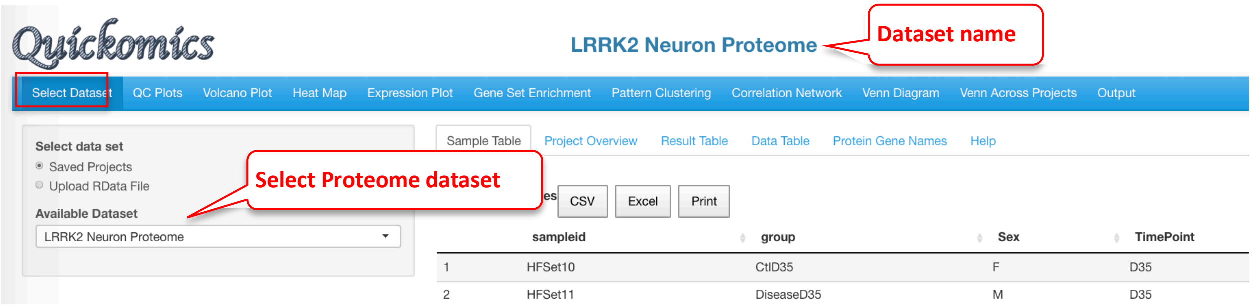

Once the dataset is selected, we go into the “Searched Expression Data” sub-tab to select a gene set.

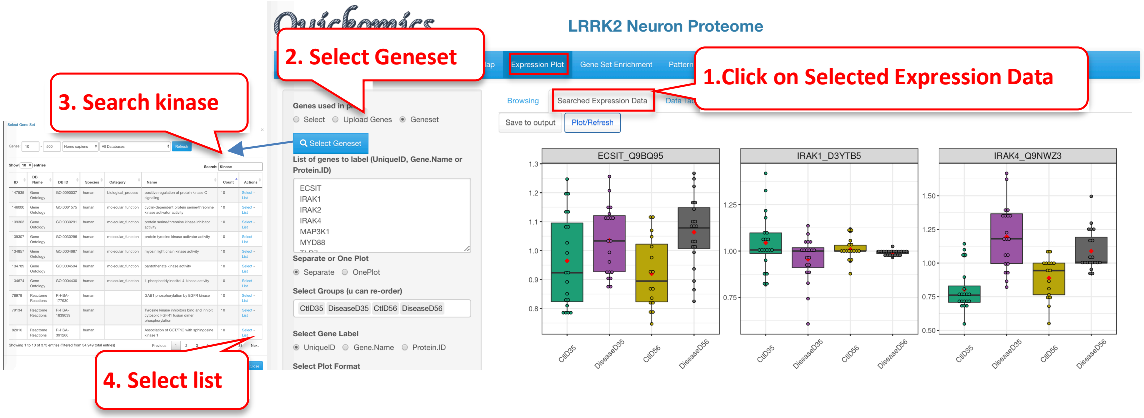

Finally, we click the “Data Table” tab to look at the expression of the protein selected for a searchable table of values.

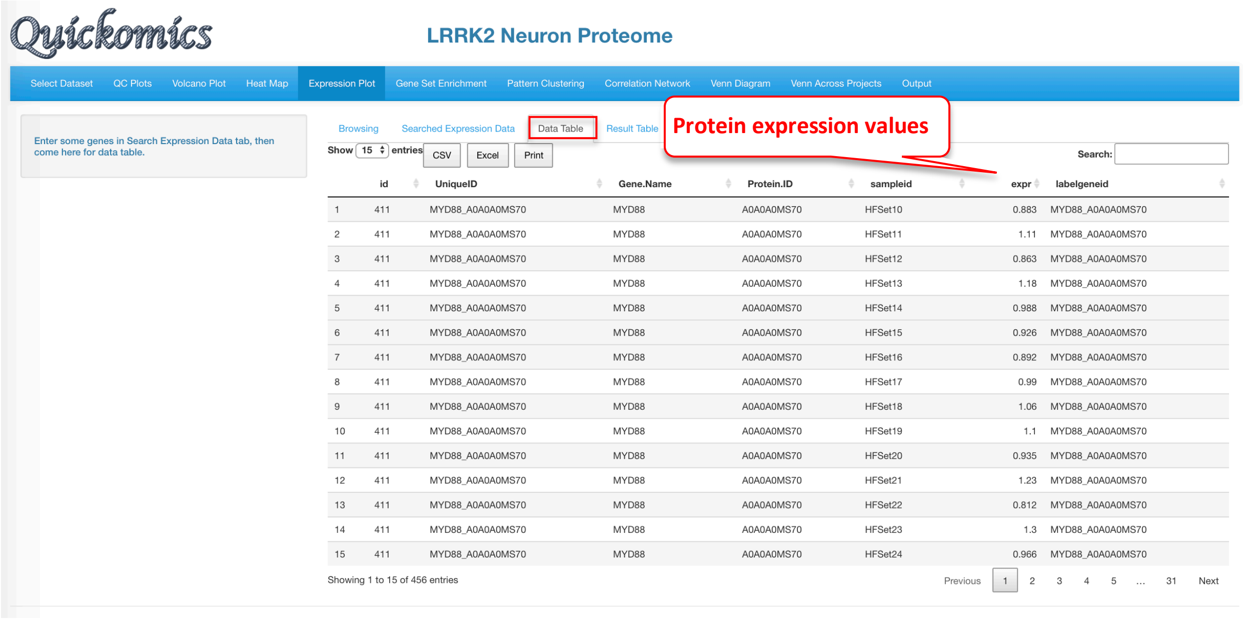

#### 6.4 Result Table

This tab lists the statistics results of the genes/proteins selected in “Searched Expression Data” as a searchable table. Each comparison defined is probed for the genes in the list and the P value and Fold changes are listed. This is specifically designed for the gene values, unlike “Data Table” in the previous section (6.3) that is designed for protein values.

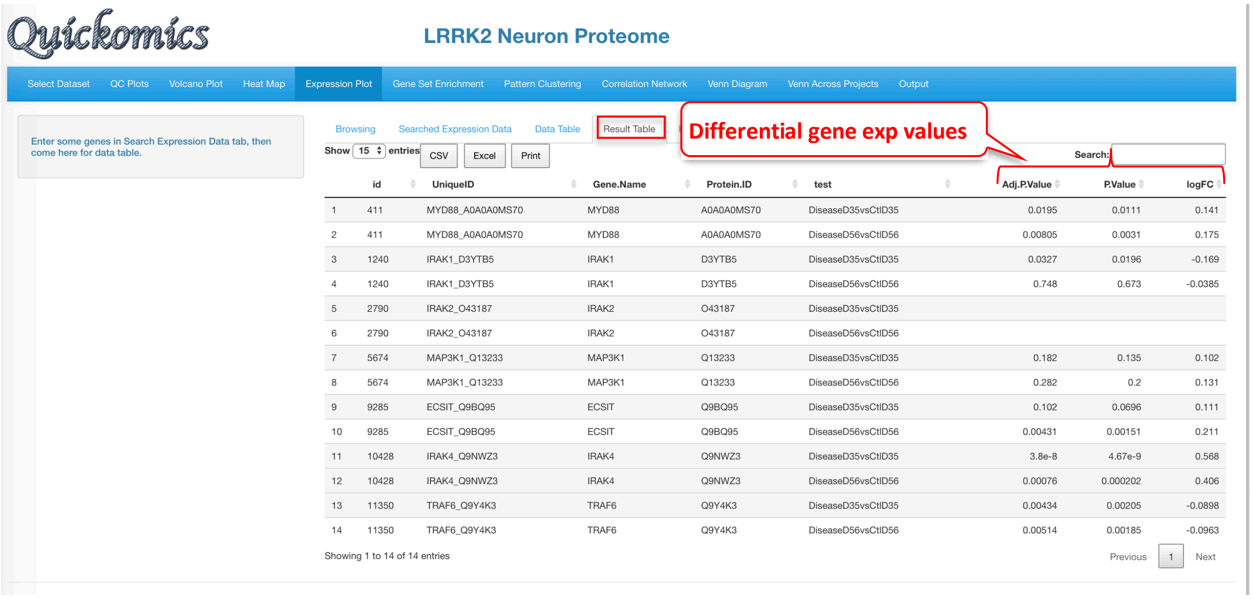

#### 6.5 Rank Abundance Curve

This functionality helps identify the relative abundance of a list of genes within a dataset. This plot helps interpret the distribution of abundance and expression levels of a set of genes. A steep gradient indicates the genes are very different in their expression levels as the high-ranking species have much higher abundances than the low-ranking species. A shallow gradient indicates that the genes have very similar expression levels as the abundances of different species are similar.

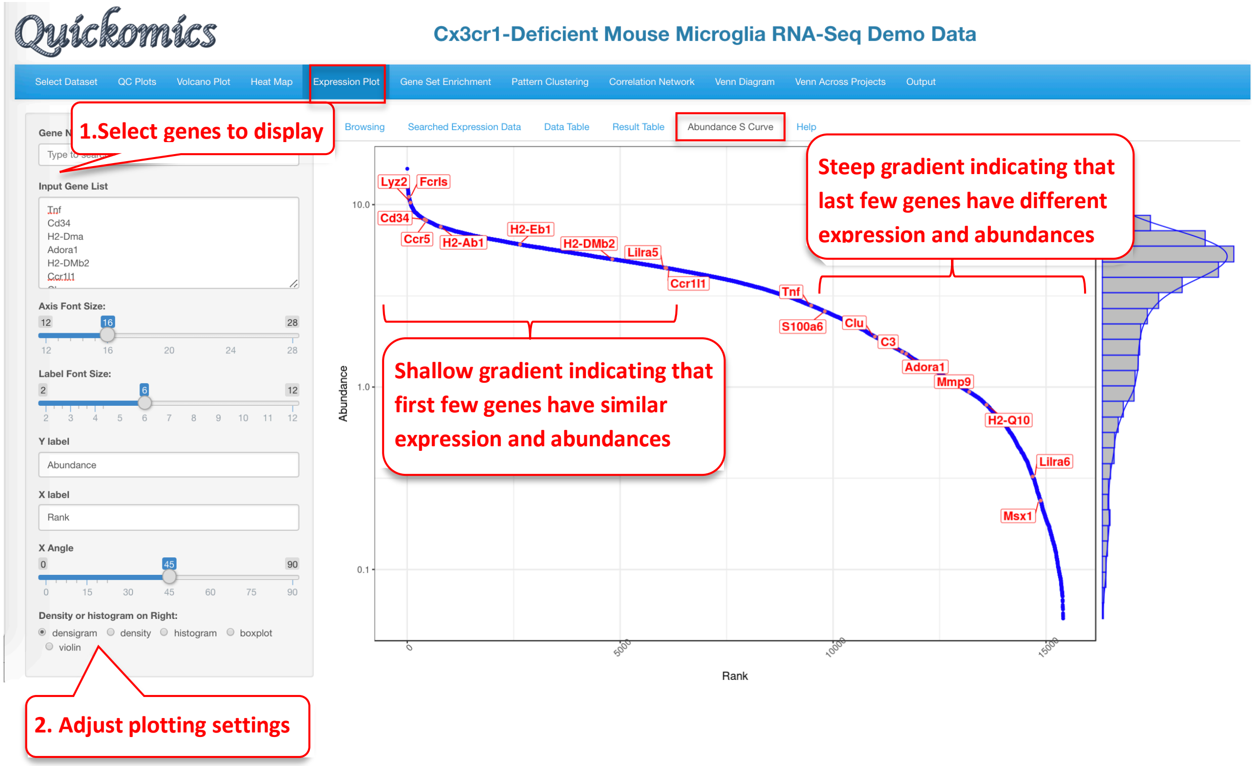

### 7 Gene Set Enrichment Module

This functionality in Quickomics is an extension to the differential gene identification. Users are able to use different cutoffs to filter out DEGs/DEPs in different comparisons and probe enriched pathways. This is a useful way to make biological sense of the data.

#### 7.1 Gene Set Enrichment

This first tab lets the users pick a particular comparison, apply different cutoff criteria and select from a list of different databases for enrichment analysis. To demonstrate usage of this functionality, we will use the Mouse Microglia RNA dataset to help answer the question which pathways were altered by Genotype and/or Age. Very similar to the results from Gyoneva et al., 2019, this analysis points to immune related terms being enriched in the down-regulated genes for 2mon_KOvs2mon_WT.

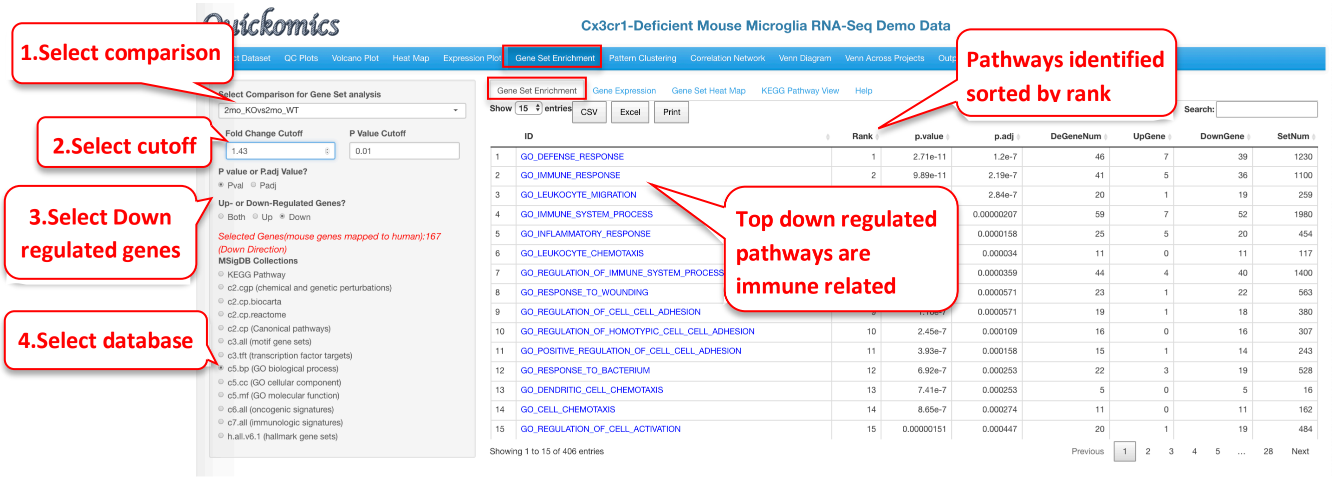

Similarly, we pull out the GO enrichment for up-regulated genes.

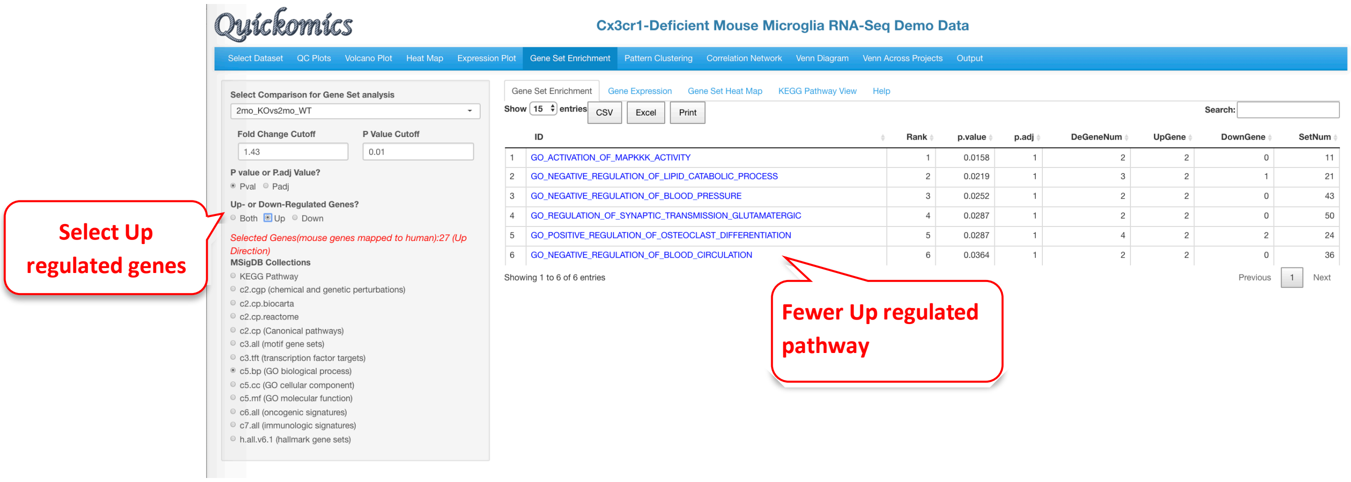

#### 7.2 Gene Expression

For the enriched terms identified in the previous tab (Gene Set Enrichment, section 7.1), Users have the ability to look at the differential values of the genes present in the set. Here we are probing the GO_IMMUNE_RESPONSE identified in the down-regulated category.

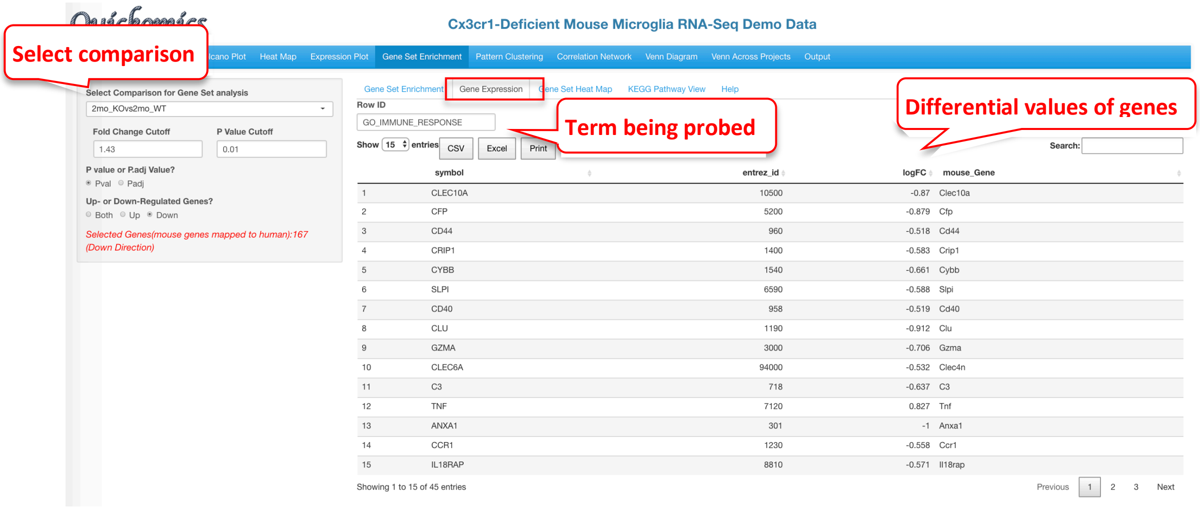

#### 7.3 Gene Set Heat Map

Like the "Gene Expression" tab, Users have the ability to plot expression of the genes in the pathway term selected as a heatmap. This visualization shows that while the Immune Response genes are enriched in the differentially expressed genes list, not all of them change in the same direction.

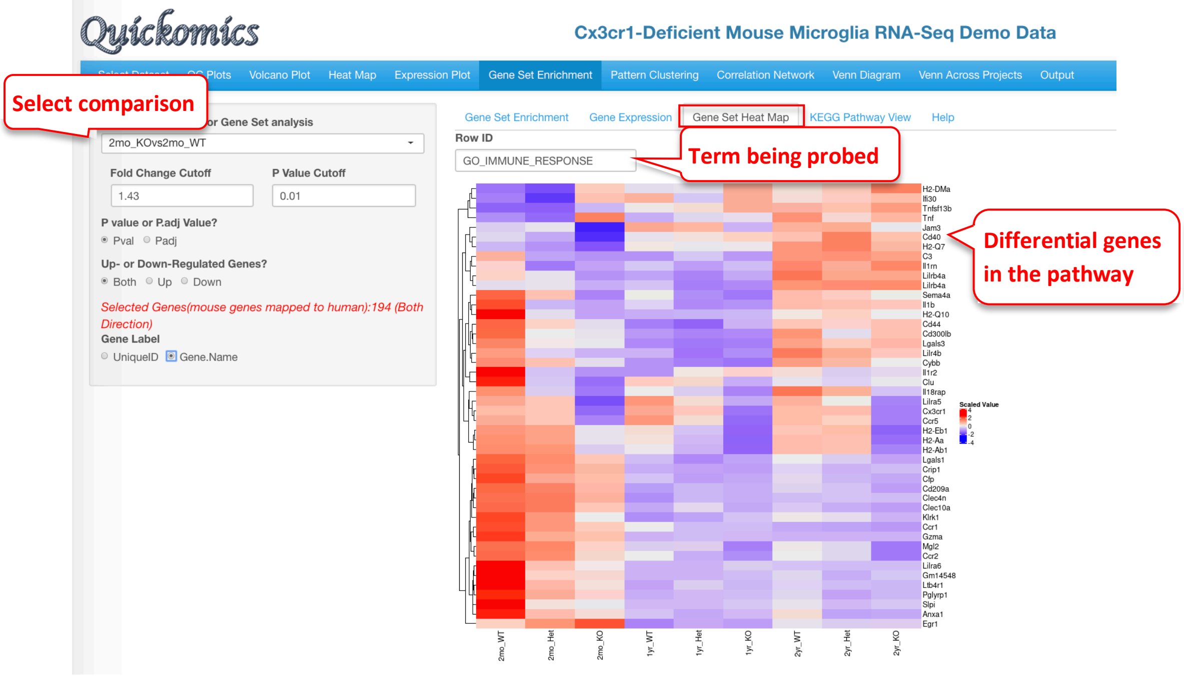

#### 7.4 KEGG Pathway View

This functionality is available only to KEGG pathway terms. Users have the option to view the Fold Change levels of different genes in the KEGG pathway selected for up to 5 comparisons. In this example we demo the KEGG pathway “hsa04612 Antigen processing and presentation” as identified as a down-regulated term in the 2mon_KOvs2mon_WT and 1yr_KOvs1yr_WT comparisons.

This figure identifies all the genes that are differentially expressed in this pathway. It is also clear that genes in the MHC I pathway do not change, but many genes in the MHC II pathway are down regulated in both the comparisons. The visualization is also useful to highlight whether or not a large proportion of genes in a pathway are altered by the perturbation.

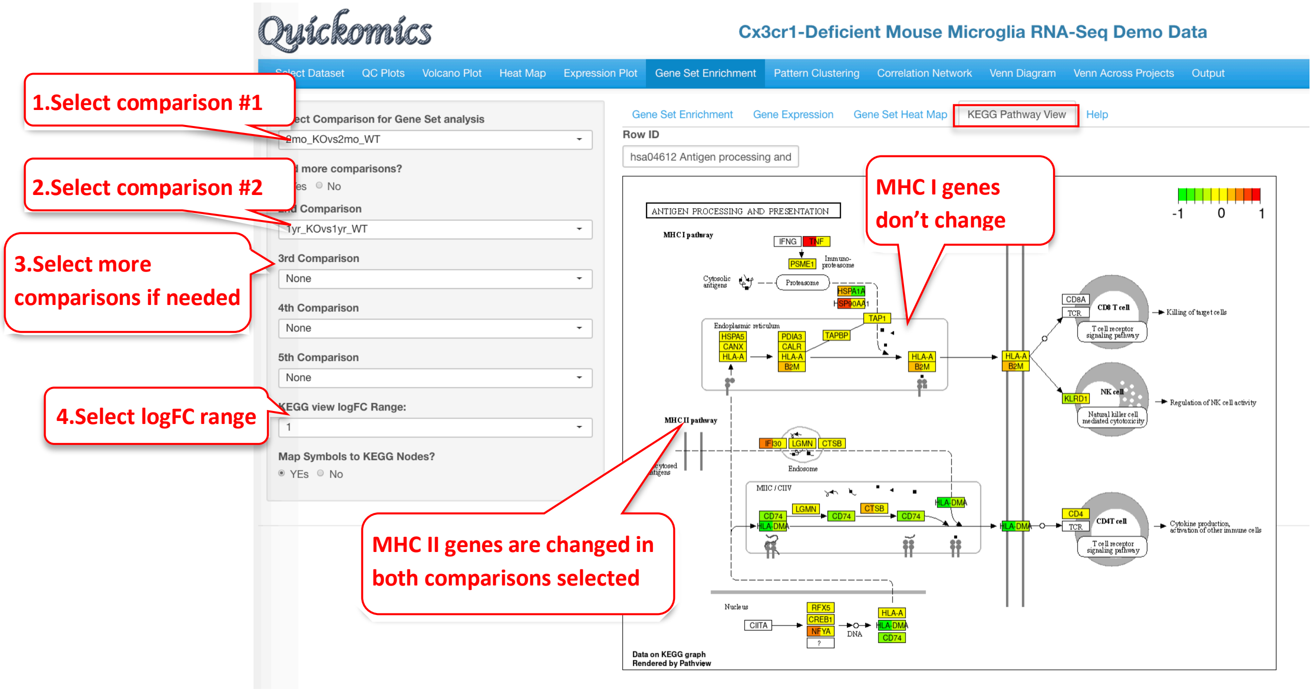

### 8 Pattern Clustering Module

This functional module in Quickomics helps cluster genes/proteins based on their expression profiles across different groups. Three different algorithms, soft (fuzzy) clustering, k-means and partitioning around medoids (PAM), are implemented.

#### 8.1 Clustering of Centroid Profiles

Users have the option to either select the DEGs/DEPS as the list of genes/proteins to cluster based off on or upload a custom gene list. DEGs/DEPs are selected based off all the comparisons, and cutoffs can be applied to limit the number of genes. In this example below, we use the top DEGs/DEPs with a Fold Change cutoff of 3 to identify 6 clusters of genes.

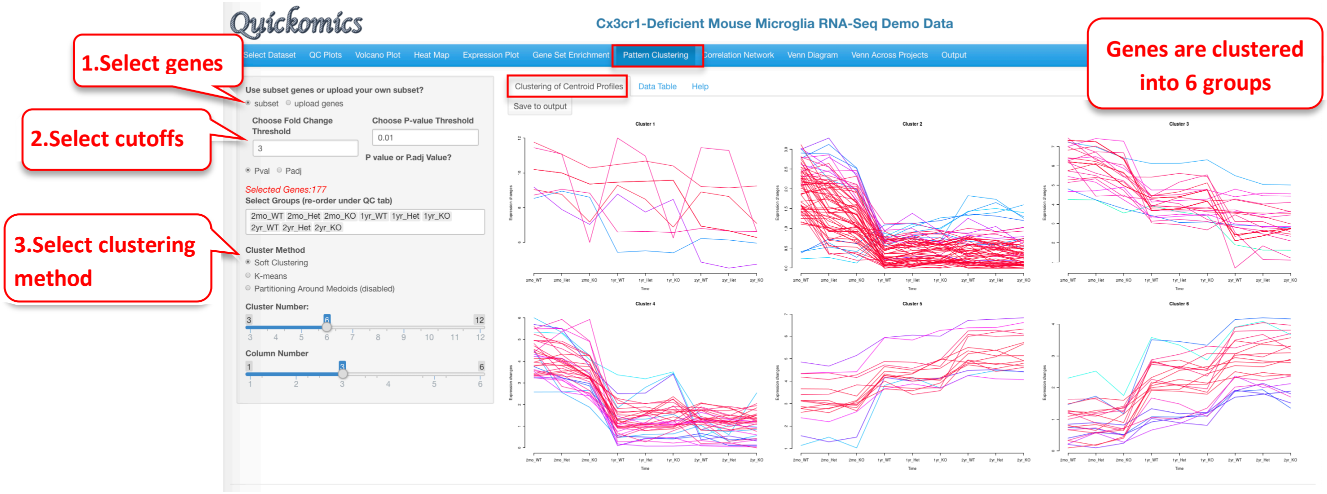

Please note that this visualization for this dataset is novel and was not previously reported in Gyoneva et al., 2019. The genes in each cluster can be viewed in the “Data Table” tab.

#### 8.2 Data Table

The data table contains genes from the previous clusters, along with their expression values

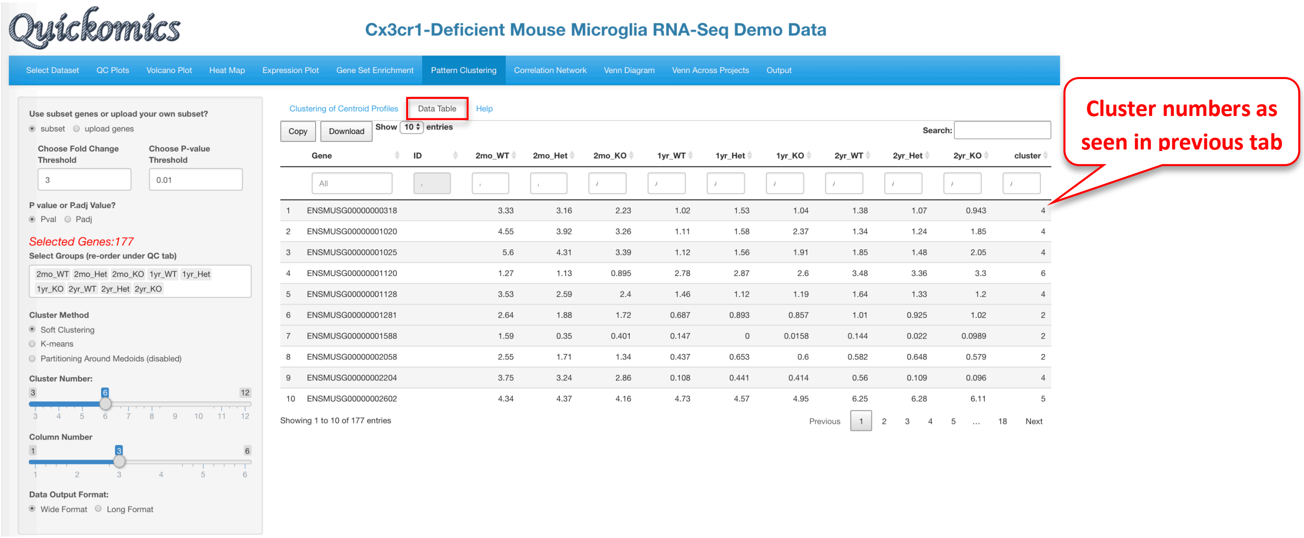

### 9 Correlation Network Module

This module helps build co-expression networks based on gene-gene or protein-protein correlations. This is especially useful during co-immunoprecipitation or pulldown proteomics experiments to identify protein partners. Users have the ability to enter 1 or more genes/proteins to identify if their expression is correlated.

#### 9.1 visNetwork

This uses the R package visNetwork to visualize the expression of correlated genes/proteins. Users have the ability to choose the R and P value cutoff for selecting the correlated genes/proteins. In this example below, we probe 3 genes and identify the expression of other genes that are correlated with them.

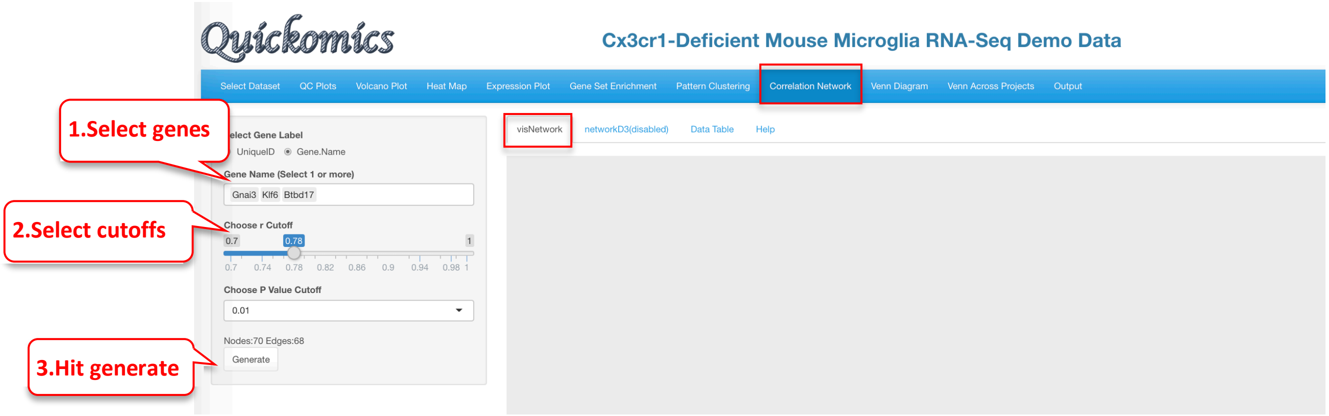

The following network plot was produced. Gnai3 is part of a large network with multiple genes correlated with it. On the other hand, Klf6 was correlated with 2 other genes, while Btbd17 was not correlated with any genes.

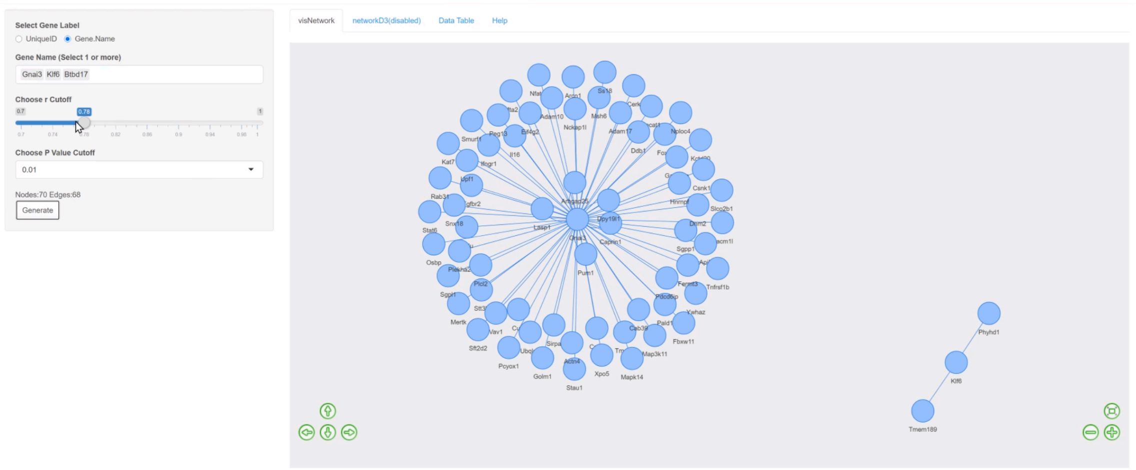

#### 9.2 Data Table

The genes identified in the previous tab as part of the network plot are available as a table in this tab. Users can perform searches and sort based on correlation statistics.

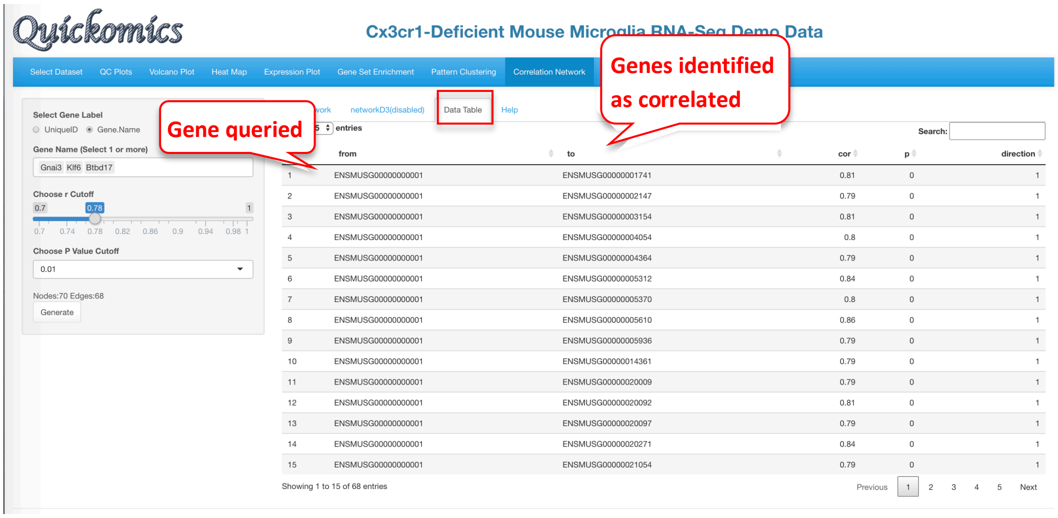
Quickomics
Cx3cr1-Deficient Mouse Microglia RNA-Seq Demo Data
Select Dataset
QC Plots
Volcano Plot
Heat Map
Expression Plot
Gene Set Enrichment
Pattern Clustering
Correlation Network
Genes identified as correlated
Gene queried
Select Gene Label
UniqueID
Gene.Name
Gene Name (Select 1 or more)
Gnai3 Klf6 Btbd17
Choose r Cutoff
0.7 0.78 1
0.7 0.74 0.78 0.82 0.86 0.9 0.94 0.98 1
Choose P Value Cutoff
0.01
Nodes:70 Edges:68
Generate
networkD3(disabled)
Data Table
Help
entries
Search:
from to cor p direction
1 ENSMUSG00000000001 ENSMUSG00000001741 0.81 0 1
2 ENSMUSG00000000001 ENSMUSG00000002147 0.79 0 1
3 ENSMUSG00000000001 ENSMUSG00000003154 0.81 0 1
4 ENSMUSG00000000001 ENSMUSG00000004054 0.8 0 1
5 ENSMUSG00000000001 ENSMUSG00000004364 0.79 0 1
6 ENSMUSG00000000001 ENSMUSG00000005312 0.84 0 1
7 ENSMUSG00000000001 ENSMUSG00000005370 0.8 0 1
8 ENSMUSG00000000001 ENSMUSG00000005610 0.86 0 1
9 ENSMUSG00000000001 ENSMUSG00000005936 0.79 0 1
10 ENSMUSG00000000001 ENSMUSG00000014361 0.79 0 1
11 ENSMUSG00000000001 ENSMUSG00000020009 0.79 0 1
12 ENSMUSG00000000001 ENSMUSG00000020092 0.81 0 1
13 ENSMUSG00000000001 ENSMUSG00000020097 0.79 0 1
14 ENSMUSG00000000001 ENSMUSG00000020271 0.84 0 1
15 ENSMUSG00000000001 ENSMUSG00000021054 0.79 0 1
Showing 1 to 15 of 68 entries
Previous 1 2 3 4 5 Next

### 10 Venn Diagram Module

This module in Quickomics helps Users identify DEGs/DEPs that are common or unique to different comparisons within one project. Users have the option to either visualize these DEGs/DEPs as a highly customizable Venn Diagram or download a table with the gene names.

#### 10.1 Venn Diagram

To view the DEGs/DEPs as a Venn Diagram, Users have the option to see the intersection of DEGs/DEPs for up to 5 different comparisons. Users can also use all DEGs/DEPs or restrict it to the ones that are Up or Down regulated.

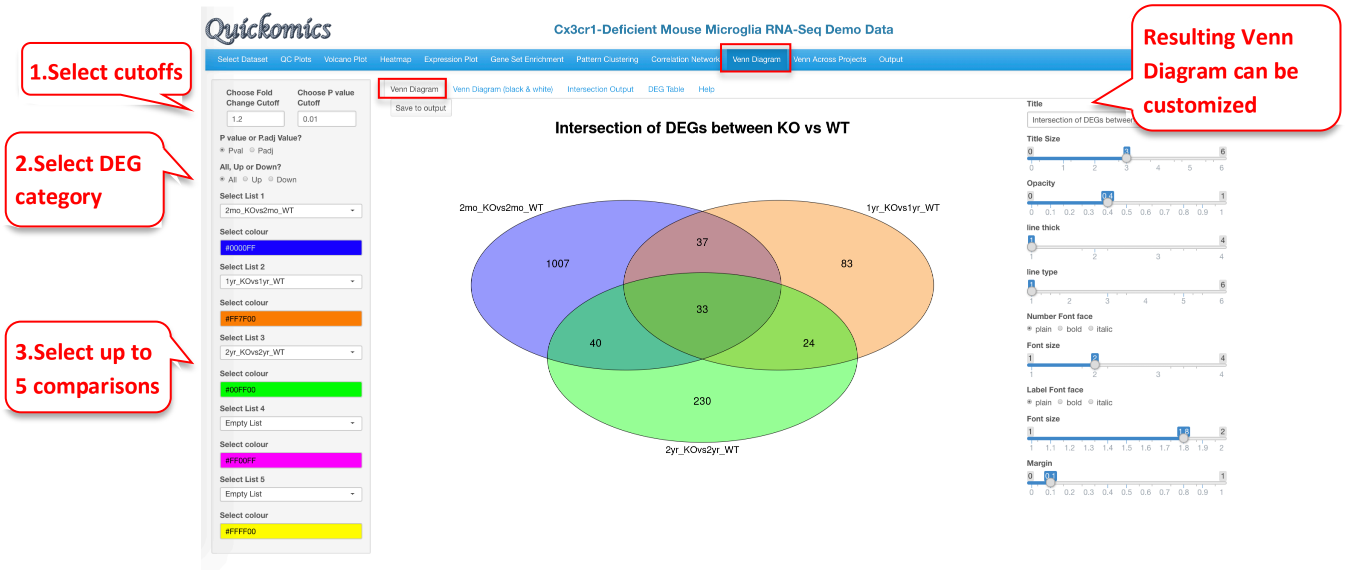

In this example there were 33 DEGs that were identified in the three comparisons probed here, which represent the set of genes altered by the KO genotype at each of the time points.

#### 10.2 Venn Diagram (black & white)

This tab has the same plot seen before in Section 10.1 but highly simplified and in black and white.

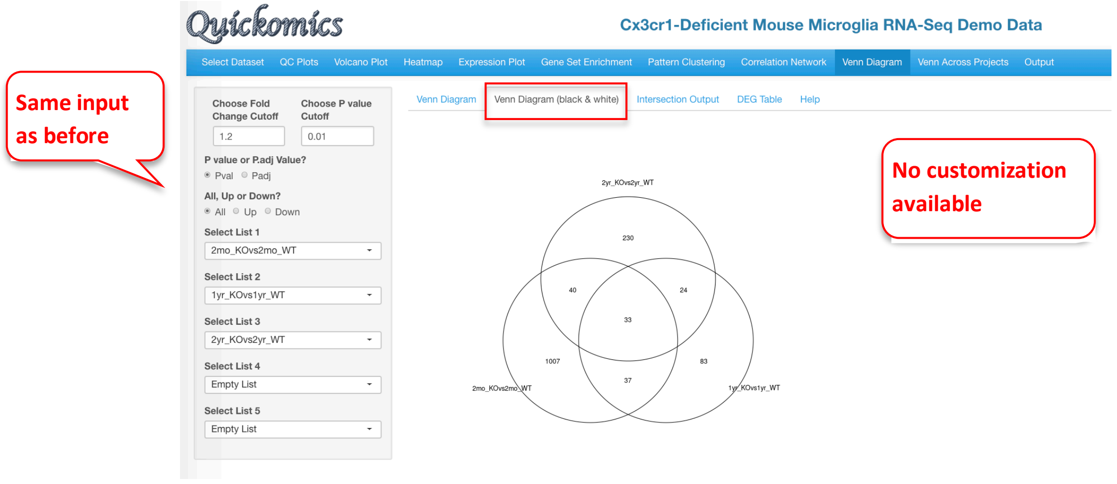

#### 10.3 Intersection Output

This tab lets the User view the individual genes that were present in the Venn Diagram and the distribution of the genes in intersection regions.

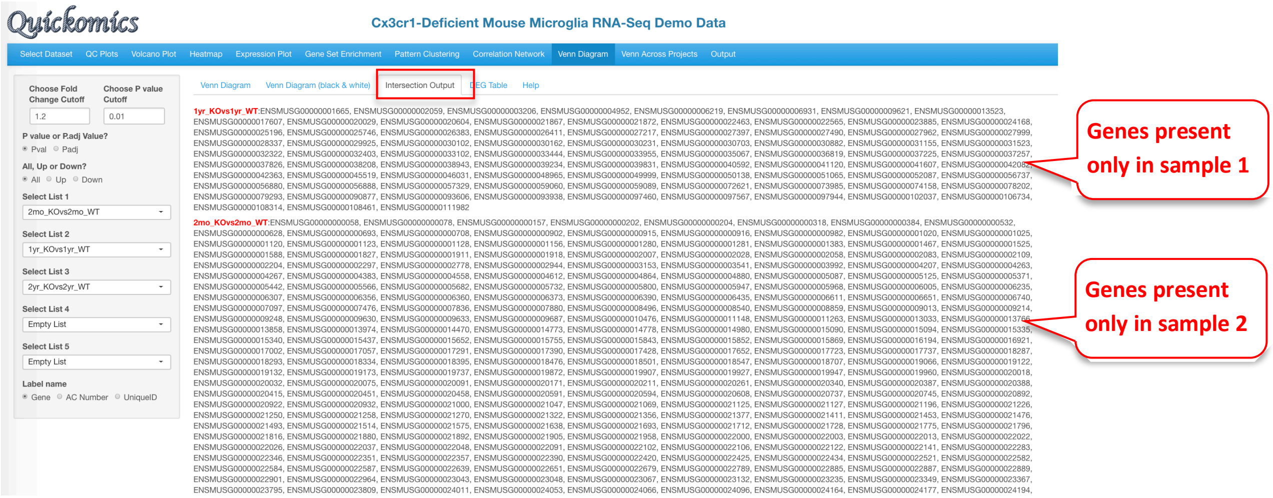

#### 10.4 DEG Table

This tab lets the Users view the Fold Change and P value of the genes identified in the Venn Diagram in the comparisons being queried.

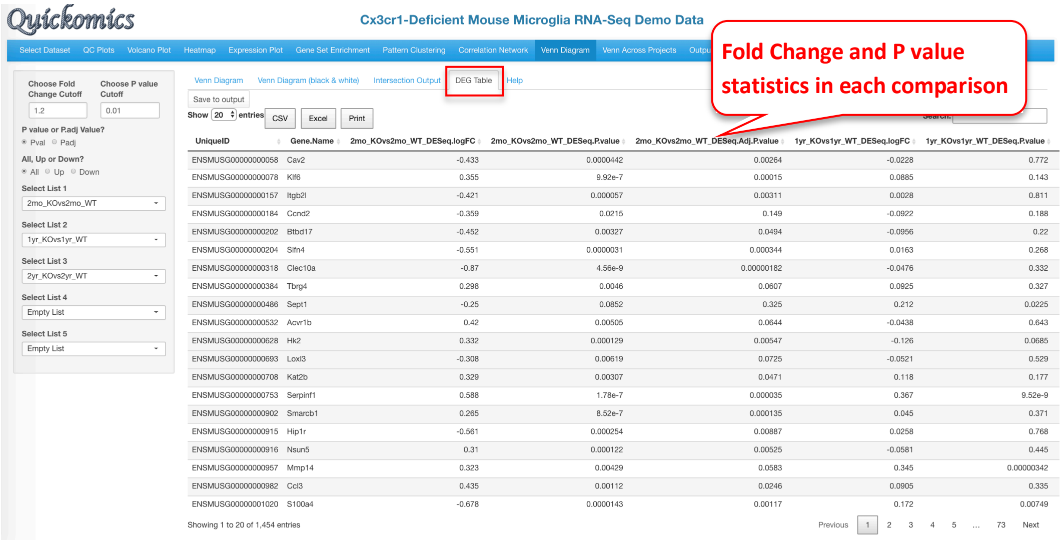

### 11 Venn Across Projects Module

This functionality in Quickomics is the only one that lets users probe more than one project/dataset together. The DEGs/DEPs identified in different comparisons across projects/datasets loaded in the interface can be compared here. By taking advantage of the “Protein Gene name” function, this module can be used to identify common DEGs/DEPs present in both RNAseq and Proteomics datasets of the same project.

#### 11.1 Venn Diagram

To illustrate this feature, we compare the two RNAseq and two Proteomic comparisons from Connor-Robson et al., 2019. We load 2 comparisons from each dataset/project and view the number of DEGs/DEPs that are common and different between the 4 comparisons.

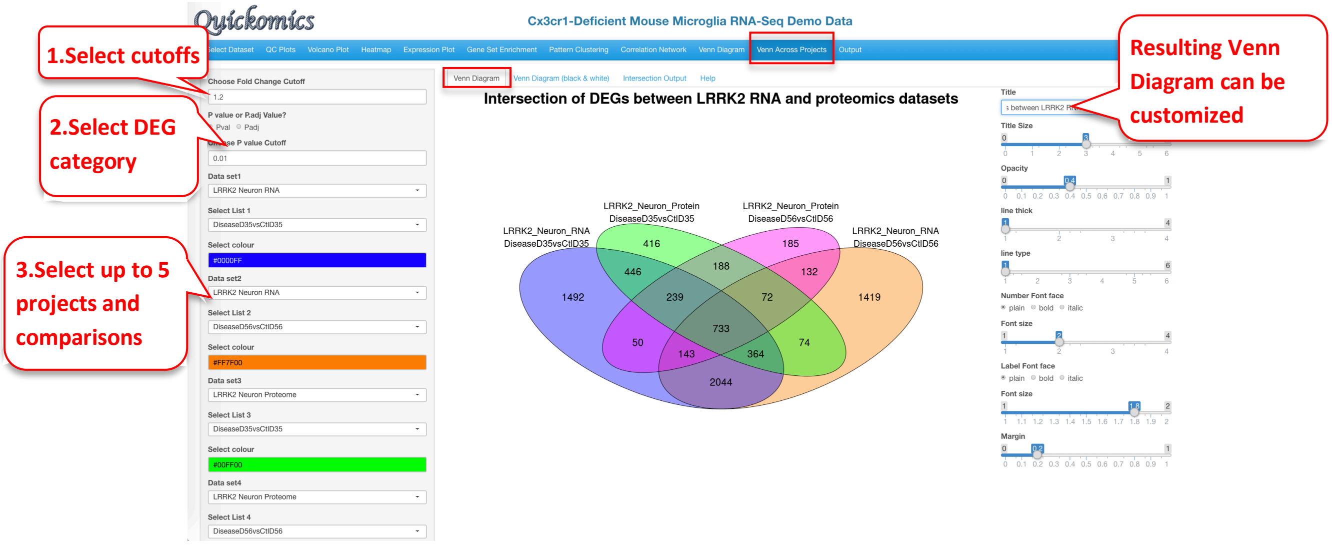

Users have the option to view this Venn Diagram as a Black & White plot.

#### 11.2 Intersection Output

Like in 10.3 Intersection Output, Users can view the individual genes present in different regions of the Venn Diagram, like the ones identified in all the samples, or the ones only present in 3 of them.

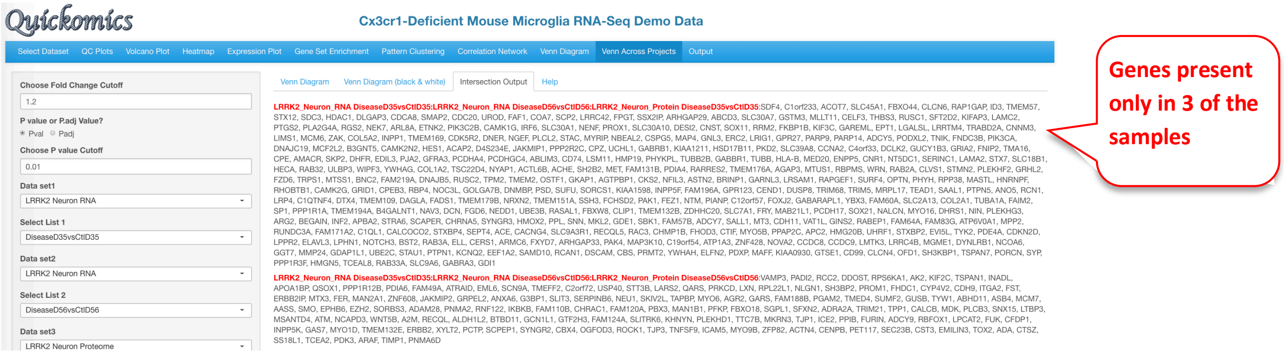

### 12 Output Module

This final tab in Quickomics is available for Users to download plots in high resolution, including for use in publications. Users can save different plots by clicking on “Save to Output” present on the top left corner of the plot during analysis and exploration of the datasets in all previous tabs. Upon clicking on the “Output” tab as seen in the image below, Users can see a list of plots that are available to download to their local machine.

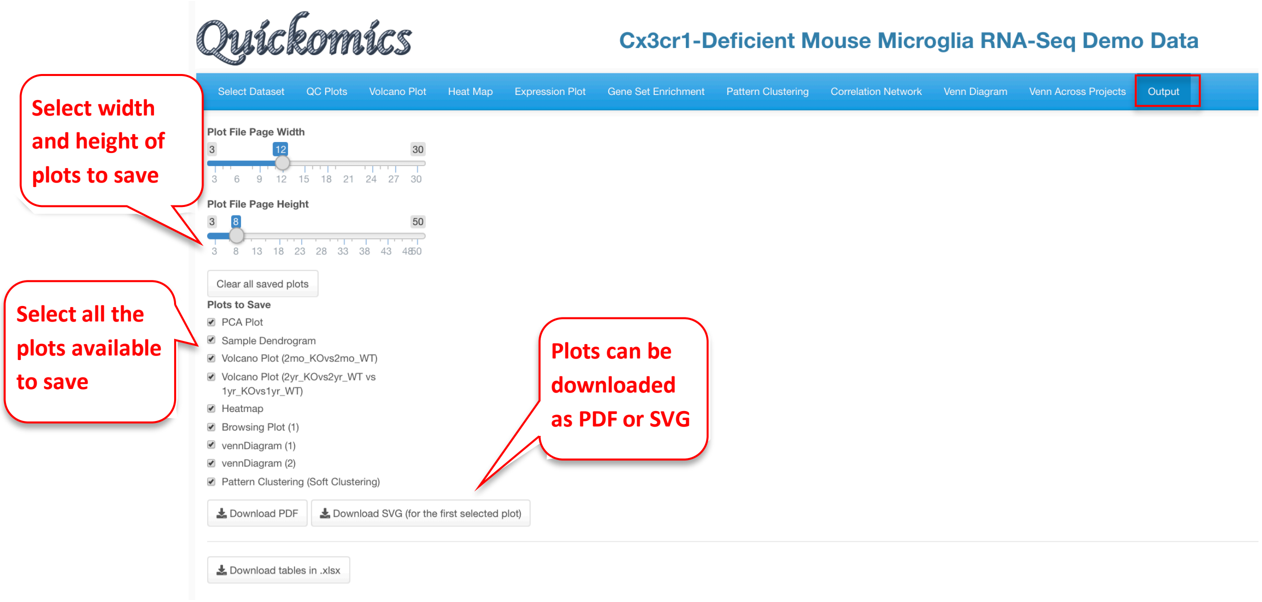
